# Mapping the Constrained Coding Regions in the human genome to their corresponding proteins

**DOI:** 10.1101/2022.09.12.507545

**Authors:** Marcia A. Hasenahuer, Alba Sanchis-Juan, Roman A. Laskowski, James A. Baker, James D. Stephenson, Christine A. Orengo, F. Lucy Raymond, Janet M. Thornton

## Abstract

Constrained Coding Regions (CCRs) in the human genome have been derived from DNA sequencing data of large cohorts of healthy control populations, available in the Genome Aggregation Database (gnomAD) [1]. They identify regions depleted of protein-changing variants and thus identify segments of the genome that have been constrained during human evolution. By mapping these DNA-defined regions from genomic coordinates onto the corresponding protein positions and combining this information with protein annotations, we have explored the distribution of CCRs and compared their co-occurrence with different protein functional features, previously annotated at the amino acid level in public databases. As expected, our results reveal that functional amino acids involved in interactions with DNA/RNA, protein-protein contacts and catalytic sites are the protein features most likely to be highly constrained for variation in the control population. More surprisingly, we also found that linear motifs, linear interacting peptides (LIPs), disorder-order transitions upon binding with other protein partners and liquid-liquid phase separating (LLPS) regions are also strongly associated with high constraint for variability. We also compared intra-species constraints in the human CCRs with inter-species conservation and functional residues to explore how such CCRs may contribute to the analysis of protein variants. As has been previously observed, CCRs are only weakly correlated with conservation, suggesting that intraspecies constraints complement interspecies conservation and can provide more information to interpret variant effects.

## Introduction

Predicting the impact of variants on protein function has traditionally been based on combining information derived from protein sequences, inter-species conservation and knowledge of the structure and function of the protein. With the emergence of multiple human genome sequences from different populations, observed variation patterns in humans provide an orthogonal source of information for assessing the impact of variants. The comprehensive catalogues of genetic variations, compiled from many human population sequencing projects, have fuelled the development of different metrics that measure the general tolerance of genes to variation. Metrics like the probability of being loss-of-function intolerant (pLI) and missense Z-scores are extensively used to prioritise genes in genome interpretation of individuals [2]. However, it is well established that ‘all parts of a protein are not equal’. The modular presence of different domains and folds can endow different regions with different functions and structural constraints [3, 4]. Hence, one would expect that regions which are extremely important for the function of a protein would be depleted of protein changing variants in healthy individuals.

In 2019 Havrilla et al. defined the Constrained Coding Regions (CCRs) as regions in the human coding genome where the following protein-changing variants were depleted: missense, stop gained, stop lost, start lost, frameshift variant, initiator codon variant, rare amino acid variant, protein altering variant, inframe insertion, inframe deletion, and splice donor variant or splice acceptor variant when affecting the protein sequence. These regions were identified in whole exome and genome sequencing data from large cohorts of healthy control populations from the Genome Aggregation Database (gnomAD2.0.1), which is currently the largest and most widely used publicly available collection of data on population variation from harmonised sequencing data [2, 5]. The premise of Havrilla et al. was that the data in gnomAD2.0.1 came from individuals who were either healthy or did not have early onset developmental abnormalities (‘healthy control populations’) and therefore their variant loci could be considered as ‘tolerated’. In the CCRs model, each of the variant-depleted (constrained) regions is weighted based on i) its length in base pairs, and ii) the fraction of individuals (above 50% of total individuals) having at least a 10x sequencing coverage at each bp of the region. A linear regression is then calculated comparing the weights and the CpG density of the regions, as an indicator of the region mutability upon spontaneous deamination of methylated cytosines. Regions with a greater weighted distance between protein-changing variants than expected based upon their CpG density (residual from the linear regression), are predicted to be under the greatest constraint. The residuals of the regression are ranked in CCRs percentiles (CCRpct) from 0 to 100, with 0 signifying unconstrained (i.e. having tolerated variants in gnomAD) and 100 being the most highly constrained regions. Put simply, the longer a constrained region and the larger its CpG content, in general, the higher its CCRpct will be. Havrilla *et al*. observed that only ∼1% of the highly constrained regions were found to be enriched with known pathogenic variants and associated with developmental disorders, and that 72% of the genes harbouring a CCR in the 99th percentile or higher were not linked yet to any disease, suggesting that CCRs could be used to reveal regions of protein coding genes that are likely to be under potentially purifying selection.

Given their relevance, it has been proposed that analysing the presence of regions like these can complement the classical procedures of phylogenetic conservation, amino acid substitution scores, and three-dimensional protein structural characterization and aid in the process of variant interpretation [1, 6]. Although recent studies have used CCRpct as an extra score for assessing the pathogenicity of variants [7–13], there has been no large-scale attempt to map the distribution of these constrained genomic regions to amino acids and to analyse their co-occurrence with different protein functional features.

The human proteome is a continuum where proteins can be fully ordered, intrinsically disordered (ID) or flexible/mobile, have a mixture of folded and ID regions or even exert transitions between both states upon binding with other proteins [14, 15]. These ID proteins and regions, can perform important and diverse functions in the cell from displaying sites for post-translational modifications (PTMs) to assembling molecular complexes that promote the phenomena of liquid-liquid phase separation (LLPS) and formation of membraneless organelles in the cell, amongst others [16–18]. Disease-causing mutations can occur in both ordered and ID regions [19, 20], and recently the focus has turned towards variants predisposing to disease in LLPS regions, mostly related to autism spectrum disorders (ASD), cancer, neurodegeneration, and infectious diseases [21–23]. However, ID regions are usually not well conserved, lack a stable protein three-dimensional structure, sequence alignments have poor accuracy and most studies and tools focus on ordered regions, making it a challenge to interpret the molecular mechanism behind disease-related variants in these regions. Hence, observing CCRs in these regions may provide some insight into their constraints during human evolution.

Herein we map the CCRs onto the protein sequence and 3D structure, by assigning the CCRpct to each amino acid site (residue) spanned by each CCR. This process has the potential to highlight key functional amino acids in both ordered and disordered proteins, lying in regions of the protein which are strongly constrained. We explore the distribution of these regions across human proteins and compare their co-occurrence with different protein functional features annotated at the amino acid level. We then perform an enrichment test of Gene Ontology (GO) terms to explore which protein classes and cellular pathways are more frequently associated with genes harbouring regions with high CCRpct.

## Results

### Mapping the CCRs to amino acids

Our aim was to explore how CCRs are distributed across the human canonical protein sequences as defined by UniProtKB/Swiss-Prot [24]. For this purpose, first, we ran the CCRs model pipeline (available in their repository: https://quinlan-lab.github.io/ccr/examples/updates) to obtain the genomic coordinates of the CCRs, but using the gnomAD3.0 dataset of variants, which aggregates 76,156 whole genomes using coordinates from the human GRCh38 genome assembly. The resulting file with the genomic coordinates of the CCRs can be obtained from our GitHub repository (https://github.com/marciaah/CCRStoAAC/blob/main/data/rawCCRs/gnomad3_0/vep101/ sort_weightedresiduals-cpg-synonymous-novariant.txt.gz). Then, we mapped the genomic coordinates of the CCRs to amino acids in UniProtKB proteins, via the Ensmebl transcripts in the GENCODE basic set (see Figure 10 A for further details), the corresponding code of the pipeline is available in our repository https://github.com/marciaah/CCRStoAAC, and the output of the mapping to amino acids is here https://github.com/marciaah/CCRStoAAC-output.

The use of GRCh38 gives a more accurate cross-mapping of genes, transcripts and proteins in the Ensembl [25] and UniProtKB databases, while using the Swiss-Prot canonical set of proteins ensures the availability of functional annotations for further analysis. From a total of 18,583 human UniProtKB/Swiss-Prot canonical proteins that matched the Ensembl protein sequences, we were able to map CCRpct to at least one region in 17,366 of them (Figure 1 A). 6,608 of the 17,366 proteins had partial coverage of CCRpct for their amino acid sites (residues) and 1,217 (from the expected 18,583) completely lacked CCRpct, as a consequence of low quality conflicted genomic regions (see Methods) that prevented the identification of CCRs. In total, about 9.8 million amino acid sites (Figure 1 B) are represented with CCRpct, out of an expected 10.7 million from the total 18,583 sequences.

**Figure 1:**
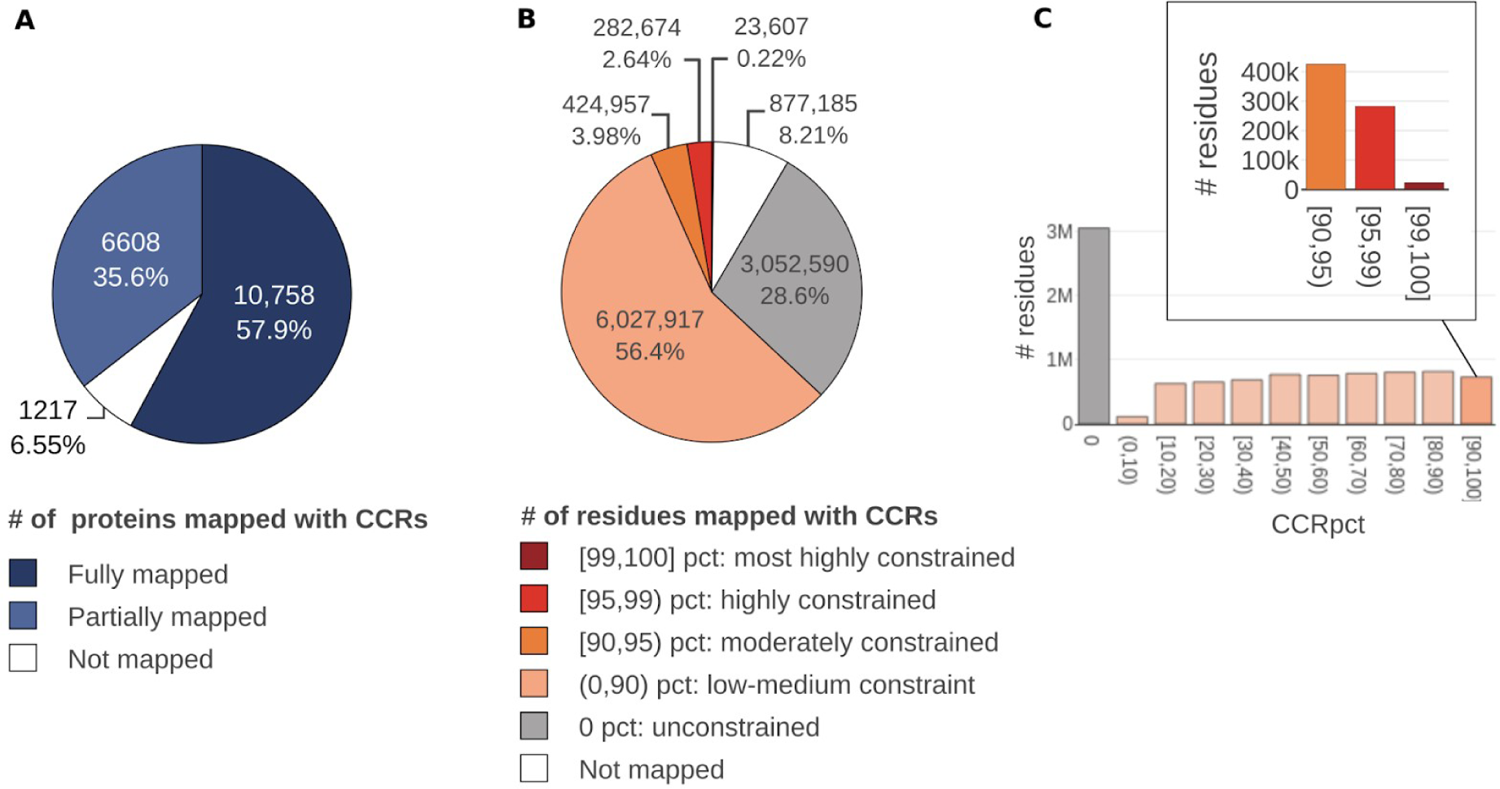
68.8% of the 9.8 million mapped amino acid sites correspond to constrained regions, ranked by different constraint percentiles. The moderately, highly, and most highly constrained positions ([90, 100] CCRpct bin) represent only 7.6% of these positions. Charts show: (A) the coverage of UniProtKB/SwissProt canonical proteins with the mapping of CCRpct, (B) the number of residues covered by the different percentiles, grouped into 5 categories, and the proportion of residues without CCRpct (not covered), and (C) the distribution of sites among the different percentiles grouped by tens, with a call-out showing the number of positions in the [90, 100] CCRpct. Interval boundary numbers should be interpreted as follows: []=included, ()=excluded, or combinations of both.

68.8% of the 9.8 million amino acids sites that we could map are in constrained regions, i.e. no protein changing variants are reported in gnomAD3.0. The remaining 3.06 million (31.2% of the mapped residues) contain at least one variant with a minimum allele count of one (Figure 1 B).

We categorised the CCRpct into different groups, as shown in Figure 1 B and C, based on the original considerations proposed by Havrilla et al, 2019, i.e., percentiles in the top 1% [99, 100] for the most highly constrained regions down to 0 for unconstrained (i.e. sites having variants in gnomAD3.0). As expected, mapping from CCRs to amino acid sites gives approximately 10% of residues in each group (Figure 1 C). The exception is for the 0-10% group, which is underpopulated at the residue level, since these CCRs are the shortest regions, with an average length of only 1.05 amino acids without variants.

To summarise, we were able to assign CCRs to 93.5% of human UniProt/SwissProt canonical proteins, equating to 91.6% of the expected residues; about two thirds of residues are constrained (i.e. without protein changing variants in gnomAD 3.0); of these sites only 0.24% residues are assigned to the top [99, 100] CCRpct bin (i.e. most highly constrained) and are exclusively from 839 regions in 751 proteins.

### Comparing CCRs percentiles with other whole gene scores (pLI and missense OEUF)

In clinical genomics and population genetics a number of metrics for assessing the overall intolerance to variability for a given gene or protein have become popular. The latest version of these scores is based on gnomAD2.1.1 [2, 26]. One, the pLI score, represents the probability of a gene being intolerant to heterozygous putative loss of function (pLoF) variants: nonsense (stop-gained), frameshift, splice acceptor, and splice donor variants. A pLI ≥ 0.9 has been used to highlight “essential” transcripts/genes/proteins. For missense and synonymous variants, Z-scores are used, measuring how far from the mean a gene is in terms of observed/expected (o/e) missense or synonymous variants. Accompanying these metrics, the authors recommend the use of upper bound fraction of the 90% confidence intervals around the o/e ratios (OEUF) for the different types of variants.

Here, we assigned pLI and OEUF scores to UniProtKB/Swiss-Prot canonical proteins with calculated CCRs, via their Ensembl transcript identifier. The mapping table, based on Ensembl (version 101) and UniProtKB (version 10-2020) can be downloaded from our repository (https://github.com/marciaah/CCRStoAAC/blob/main/data/mapping_tables/ ensembl_uniprot_MANE_metrics_07102020.tsv.gz). For our analysis, we used the recommended thresholds: pLI ≥ 0.9 and missense OEUF ≤ 0.35, as a simple way to define a gene/protein as highly constrained for pLoF and missense variants, respectively.

We observed 2,916 ‘essential’ proteins with pLI ≥ 0.9, and 75% of these have at least one region that scores with very high CCRpct in the range [95, 100]. Also, only 113 proteins presented missense OEUF ≤ 0.35, and 93% of them have CCRpct in the [95, 100] group. However, when we looked at proteins with pLI < 0.9 or OEUF > 0.35, we found that about a quarter or more of these more “variant-tolerant” proteins also include highly constrained regions, which are distributed across the proteins with all values of missense OEUF and pLI (Supplementary Figure 1). These observations highlight the importance of looking at local constraint scores, using CCRpct, in order to understand more deeply the impact of variants in protein coding genes.

### The correlation between CCRs percentiles, interspecies conservation and length of the regions

We next explored how the CCRpct (based on intra-human variation) correlates with interspecies amino acid conservation for each amino acid position in the human proteome. Figure 2 A shows that the average interspecies conservation increases with increasing CCRs percentile. However, there is a surprisingly large variability within each percentile category (Figure 2 A and B) and the overall correlation is very low (Pearson=0.11). This agrees with the previous observations (Havrilla et al. 2019) that the correlation between the CCRpct and the average GERP++ nucleotide conservation scores [27] for the regions is very low and hence the intra-species conservation in humans complements the interspecies conservation.

**Figure 2:**
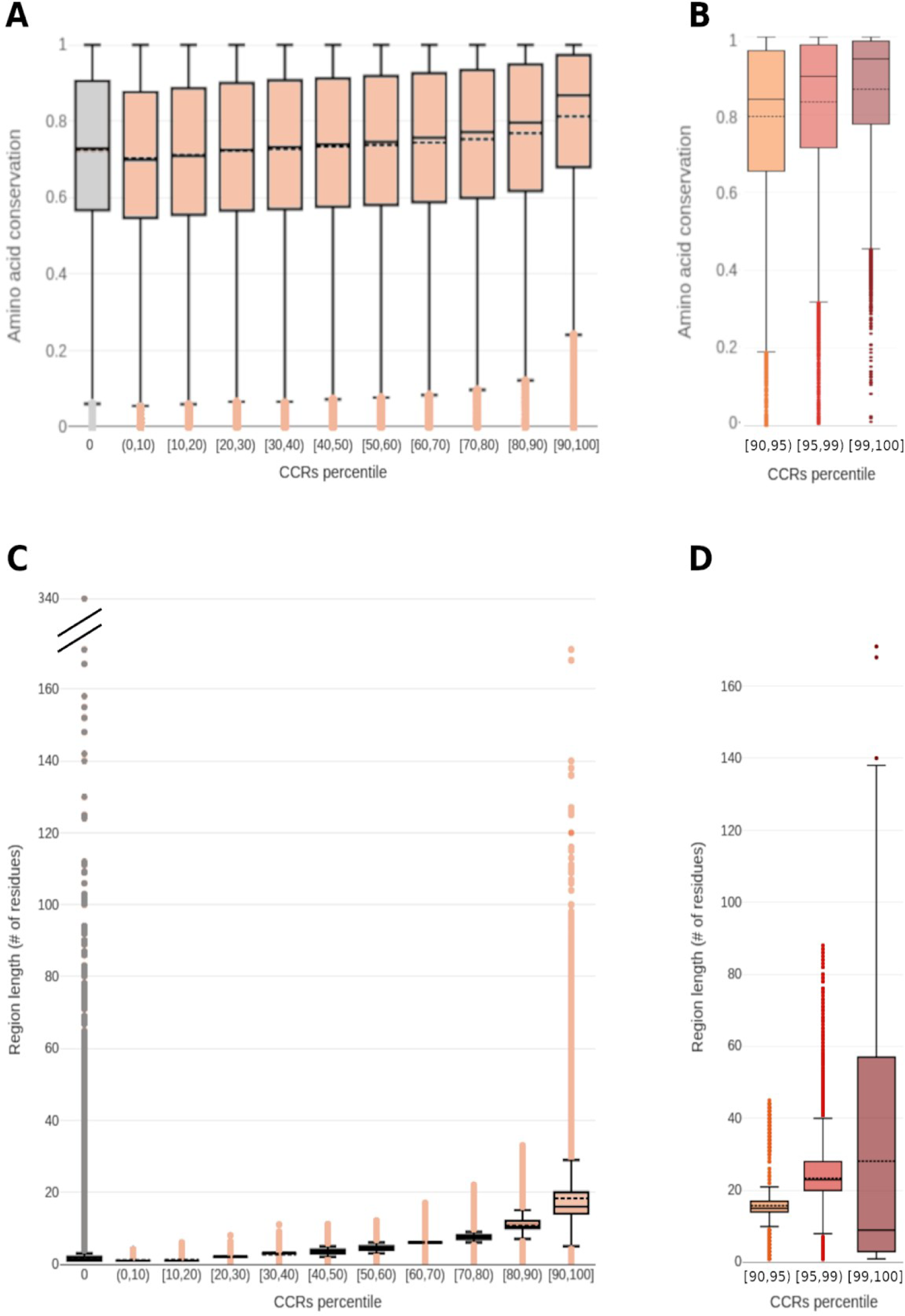
The higher the constrained percentile the larger the mean length of the regions and the higher the mean conservation of the amino acids within them. However, each percentile category exhibits a high variability. The boxplots in the upper panels, (A) and (B), show the distributions of amino acid conservation for all the mapped positions. In the lower panels, (C) and (D), the distribution of lengths for all the mapped regions are shown. Left panels (A) and (C) show all the CCRpct for unconstrained (0 pct) and constrained categories, sub-categorised into intervals of ten. The right panels, (B) and (D), focus only on the distribution of the subset of medium to highly constrained positions and regions, i.e. percentiles [90, 100], subdivided into three categories from medium to most highly constrained. The colouring follows the same criteria as described in Figure 1 B.

In a similar way, for all amino acid positions we compared the length of the CCRs (in number of amino acids) against their percentiles (Figure 2 C and D). The average length of regions increases with CCRs percentile, which is expected given that CCRs are prioritised by region length. Nevertheless, there is a high variability within each CCRs percentile category. The most highly constrained regions (percentiles [99, 100]) are only present in proteins of at least 100 amino acids in length (Supplementary figure 2).

To explore the numbers of amino acid sites having different combinations of CCRpct and conservation scores, we stratified both measures into ten groups and built a 2D matrix counting the numbers of residues in each cell. The resulting distributions for constrained and unconstrained sites are shown in Figure 3. The majority of both constrained and unconstrained positions have conservation scores >0.4 and distribute evenly in a plateau up to a conservation of 0.95. Above this level, the counts increase, in particular for the two extremes of more constrained (percentiles [90, 100]) and unconstrained sites (percentile 0). The 3D surface shown in Figure 3 highlights the disparity between these CCRpct and conservation scores and the high frequency (importance) of residues which are completely conserved (ScoreCons=1)[28]. CCRpct are able to differentiate between such residues, according to observed variation and length of conserved regions, providing a valuable score for analysis.

**Figure 3:**
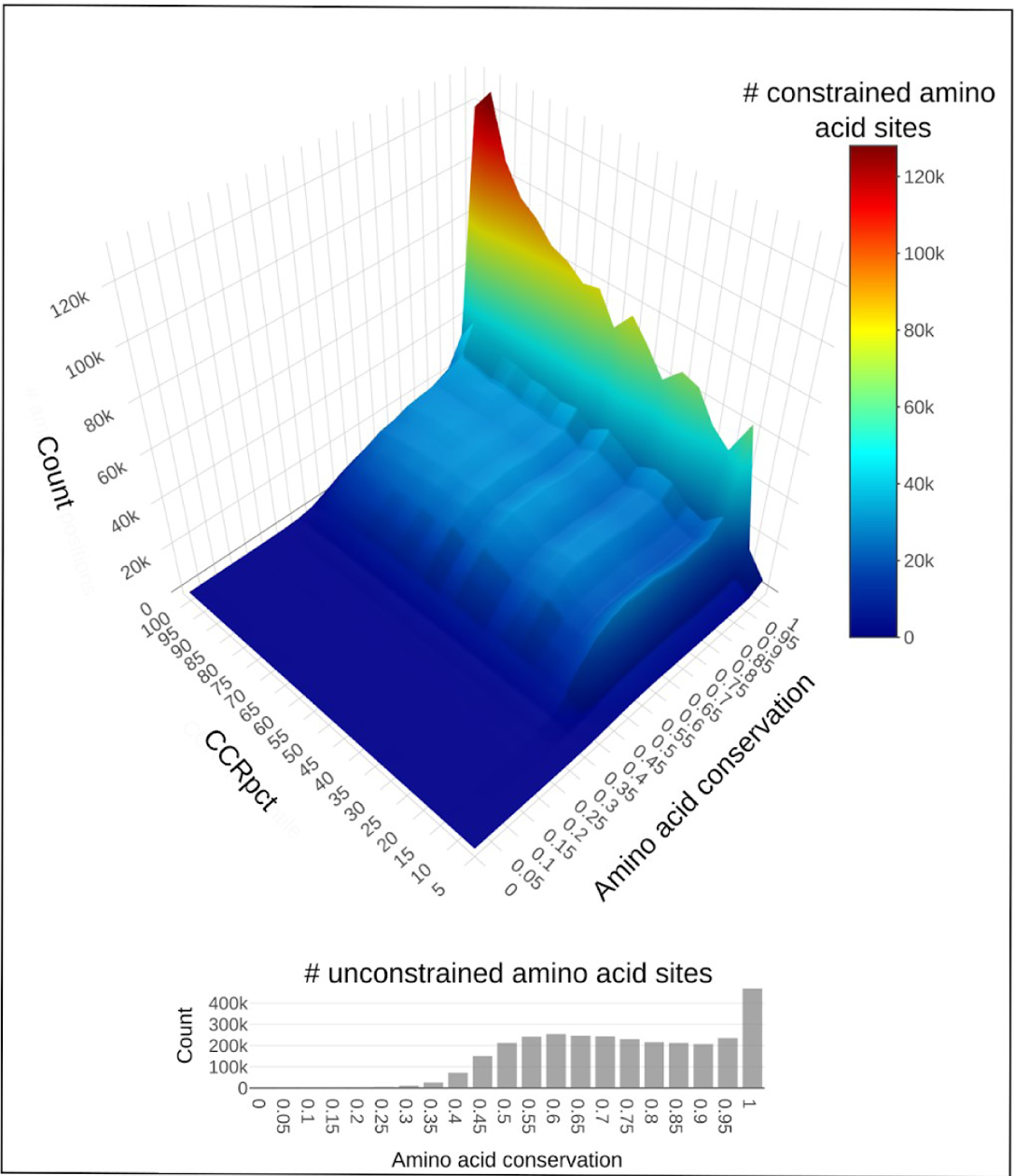
Distribution of counts of amino acid sites in different bins of CCRpct and conservation scores. The 3D heat map shows counts for constrained amino acid sites, while the histogram shows the unconstrained ones.

### Protein features and CCRs percentiles

Protein annotations from UniProtKB/Swiss-Prot [24], PDBe [24, 29], VarSite [30], M-CSA [31], BioLip [32], MobiDB [33], ELM [34], Ensembl [35] and ClinVar [36] databases were obtained and aggregated for each amino acid site in the human canonical and annotated proteins of UniProtKB. 9.8 million sites were annotated in this way, assigning CCRpct and the 30 protein annotations listed in Figure 10 (Methods).

Figure 4A and B presents a broad overview of the distribution of total sites annotated with different protein features, with the corresponding distributions of conservation scores and length of regions for such sites. For simplicity, we only present this information for the two extremes of CCRpct: the more constrained sites in percentiles [90, 100] and the variable or unconstrained sites with percentile 0.

**Figure 4:**
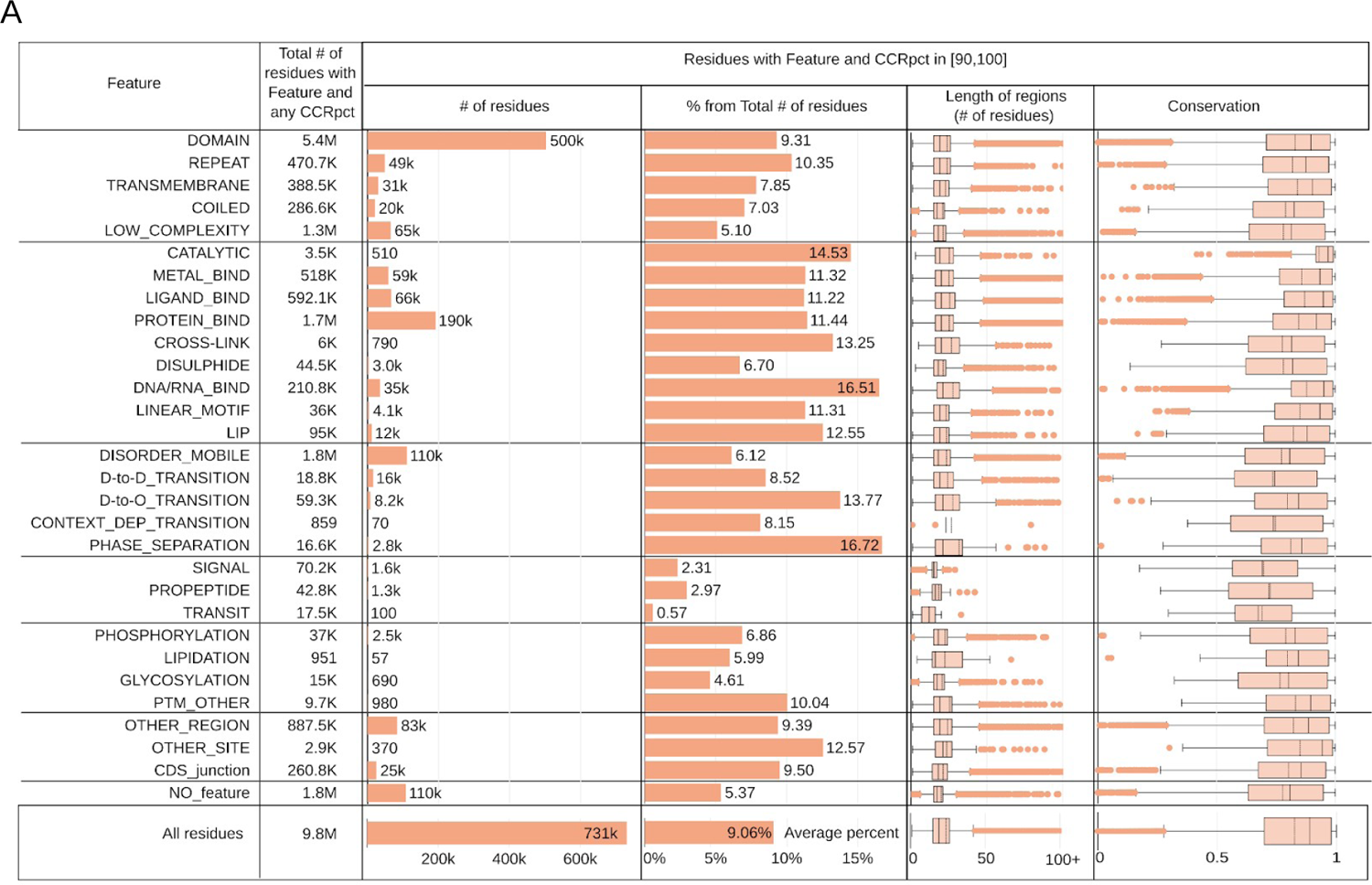
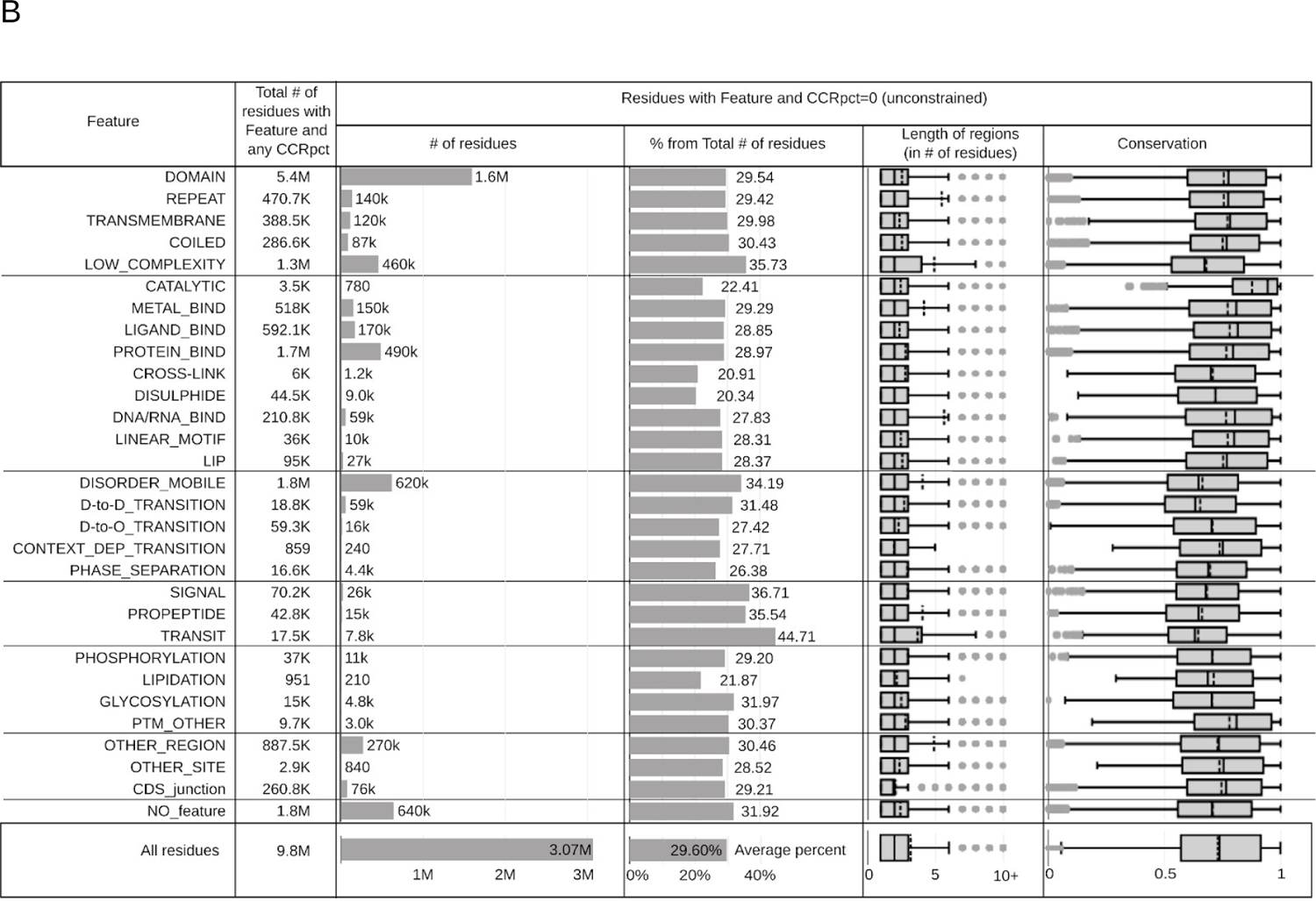
Distribution of amino acid sites corresponding to A) highly constrained regions with percentiles in the interval [90, 100] and B) unconstrained regions harbouring tolerated variants in gnomAD3.0, and in coincidence with the different protein features as listed in the first column. “All sites” panels at the bottom correspond to counting all the sites without distinction of protein features. Average percent= average of all the % from Total # of residues.

In order to investigate in detail how different protein features are constrained across human populations, we calculated odds ratios (OR) to measure the enrichment of residues in CCRpct categorised in 7 groups for each of the 30 protein feature annotations, compared to a random distribution (see in Methods, *Odds ratios tests for enrichment: I. CCRpct and presence of each one of the 30 protein features*). Figure 5 presents forest plots for comparing the resulting OR (listed in Supplementary Table 1).

**Figure 5:**
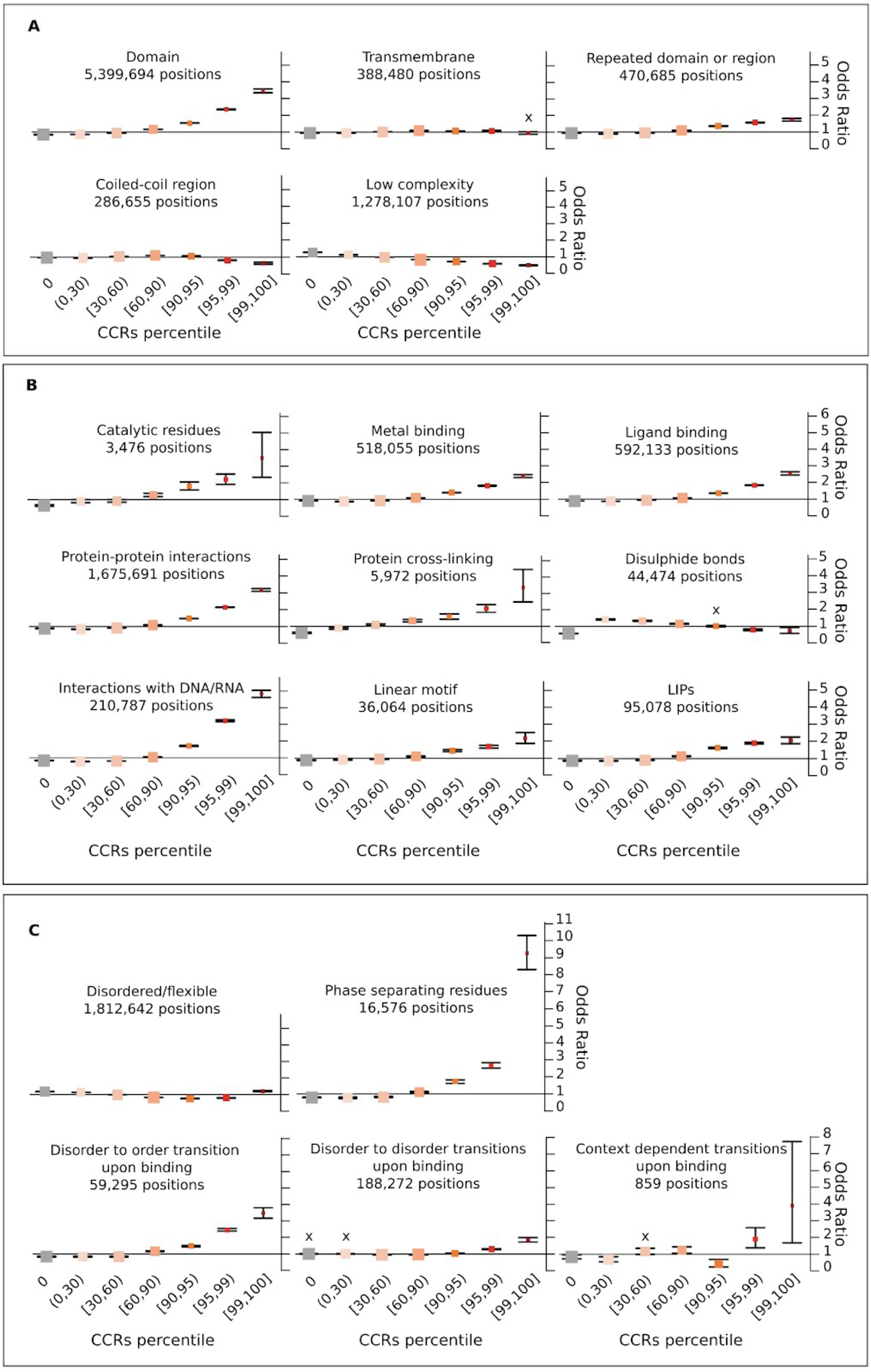
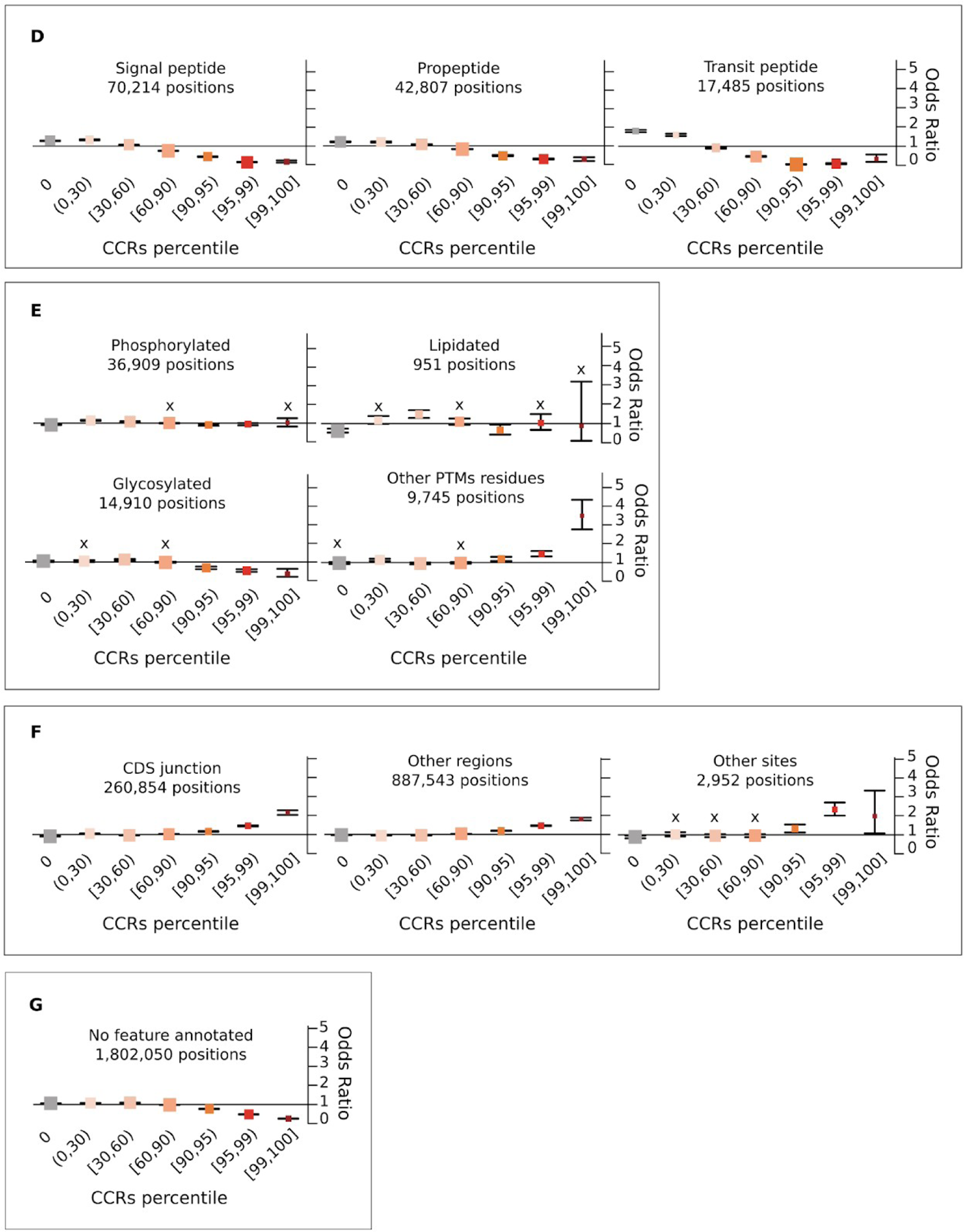
Propensity of co-occurrence of amino acid positions in specific protein features or functional sites with the different CCRpct. Odds ratios (OR) forest plots with 95% confidence intervals (CI) based on two-tailed Fisher’s exact test for amino acid sites in (A) domains, globular, non-globular and compositionally-biased protein regions, (B) interactions and catalysis, (C) disordered/mobile residues, structural transitions between disorder/flexibility and order upon binding and regions driving LLPS, (D) different signalling regions, (E) post translationally modified sites, (F) coding DNA sequence (CDS) junctions and other functionally relevant regions annotated in UniProtKB, and (G) residues lacking any of the features or functional annotation considered in this work. The vertical lines through the boxes illustrate the length of the CI. The line at OR=1 is the line of no clear difference, boxes and intervals above this represent co-occurrences more likely to happen, while boxes and intervals under the line represent the contrary. A cross (X) above a box depicts an OR where the association is statistically not significant (i.e. p-value > 0.05 or the 95% CI crosses over OR=1). See Supplementary Table 1 for all the p-values corresponding to the two and one tailed Fisher’s exact tests.

Additionally, we performed OR tests for assessing the overall enrichment of sites with the different protein features and their co-occurrence with inter-species conservation scores and CCRpct. We did this by defining 6 different groups, as described in Methods, *Odds ratios tests for enrichment: II. CCRpct and conservation with the presence of protein features*. Put simply, we divided the cells of the heatmap in Figure 3 into 4 quadrants and the histogram for unconstrained sites into 2 halves, counted the numbers of residues and calculated the ORs. Supplementary Table 2 shows the resulting OR and Table 1 summarises the features enriched in each group combining CCRpc and conservation score.

**Table 1:** List of features that are more enriched in the different combinations of CCRpct (row-wise) and conservation (column-wise), sorted by OR in each cell (only showing OR≥1.1). OR and p-values are between parenthesis, with stars depicting significative Fisher p-values: ‘*’ ≤ 0.05, ‘**’ ≤ 0.01, ‘***’ ≤0.001. The full list of features and values is in Supplementary Table 2. In bold we highlight the most enriched features, with OR≥1.5.

|  | Lower conservation<br>(ScoreCons ≤0.5) | Higher conservation<br>(ScoreCons >0.5) |
| --- | --- | --- |
| Higher CCRpct<br>(50,100] | **D_to_D (1,90\*\*\*)**<br>**PROPEP (1,50\*\*\*)**<br>DISORDER_MOBILE (1,46***)<br>CONTEXT_DEP (1,45*)<br>D_to_O (1,44***)<br>PHOSPHO (1,25***)<br>NO_feature (1,15***)<br>LLPS (1,13***) | **CATALYTIC (1,90\*\*\*)**<br>**CROSSLINK (1,63\*\*\*)**<br>**LLPS (1,50\*\*\*)**<br>DNA/RNA (1,46***)<br>DOMAIN (1,42***)<br>DISULPHIDE (1,35***)<br>MOTIF (1,34***)<br>LIP (1,33***)<br>PROTEIN (1,32***)<br>LIGAND (1,31***)<br>D_to_O (1,30***)<br>METAL (1,28***)<br>REPEAT (1,25***)<br>CONTEXT_DEP (1,25**)<br>SITE (1,22***)<br>TRANSMEM (1,20***)<br>LIPID (1,19**)<br>CDSjunction (1,16***)<br>PTM_OTHER (1,12***)<br>REGION (1,11***)<br>COILED (1,10***) |
| Lower CCRpct<br>(0,50] | **TRANSIT (1,83\*\*\*)**<br>**PROPEP (1,79\*\*\*)**<br>**D_to_D (1,71\*\*\*)**<br>**DISORDER_MOBILE (1,62\*\*\*)**<br>LOW_COMPLEXITY (1,39***)<br>PHOSPHO (1,22***)<br>NO_feature (1,17***)<br>SIGNAL (1,11***) | **DISULPHIDE (1,74\*\*\*)**<br>**LIPID (1,56\*\*\*)**<br>SIGNAL (1,27***)<br>TRANSIT (1,16***)<br>CARBOHYD (1,15***)<br>PTM_OTHER (1,14***)<br>CDSjunction (1,10***) |
| Unconstrained<br>CCRpct [0] | **TRANSIT (1,86\*\*\*)**<br>**PROPEP (1,81\*\*\*)**<br>**D_to_D (1,69\*\*\*)**<br>**DISORDER_MOBILE (1,67\*\*\*)**<br>**LOW_COMPLEXITY (1,61\*\*\*)**<br>SIGNAL (1,23***)<br>NO_feature (1,10***) | **TRANSIT (1,58\*\*\*)**<br>SIGNAL (1,24***)<br>LOW_COMPLEXITY (1,14***)<br>CARBOHYD (1,11***) |

We grouped the different features for discussing their enrichments with CCRpct and with CCRpct and conservation, as is summarised below:

I. ***Domains and compositionally-biassed protein regions.*** This group includes large features (e.g structural domains) and biassed sequences (e.g. coiled-coils) (Figure 5 A). The most striking ORs distribution occurs for domain regions - (ie those regions of a protein classified as being in a domain, according to Pfam or CATH) compared to those lying outside such a domain. There is a clear indication that amino acid sites within domains are more constrained than those outside. The repeated domains show a similar, but weaker correlation. Surprisingly, the transmembrane regions showed little if any enrichment for highly constrained regions, perhaps reflecting the lipid environment where variants between the hydrophobic amino acids are common. The residues in low complexity regions are preferentially unconstrained and rarely show the highest levels of constraint.
II. ***Interactions and catalytic residues.*** Residues involved in catalytic sites, binding to metals and/or ligands, protein-protein interactions, protein-protein cross-linking, interactions with DNA/RNA, linear motifs, and linear Interacting peptides (LIPs) are all more likely to be associated with medium to high percentiles of constraint (in the range [60, 100]) (Figure 5 B). Most of these residues are also less likely to be associated with unconstrained regions, i.e. presence of gnomAD tolerated variants. Catalytic sites, in particular, presented the highest odds of having high CCRpct and high conservation (OR=1.9, in Table 1), and the average residue conservation is consistently high in combination with all the CCRpct, including the unconstrained sites where gnomAD tolerated variants are located (Supplementary Figure 5). Disulphide bonds are an interesting exception - showing no preference to lie in a highly constrained region, but also they are rarely unconstrained. In particular, catalytic sites, disulphide bonds and cross-linking covalent linkages have ORs that suggest they are of the order of 0.5-0.6 times as likely to have tolerated variants, while for sites involved in other interactions the ORs are between 0.7 and 0.91.
III. ***Disorder related features.*** The 1.8M amino acids that were annotated in our dataset as intrinsically disordered (ID) or mobile and with CCRpct assigned (about 18.5% from the total of 9.8M residues) showed two contrasting tendencies. These sites are more likely to coincide with unconstrained and very lowly constrained sites (percentiles [0,30)) and also with the top most highly constrained percentiles [99, 100] (Figure 5 C). ID regions/proteins tend to evolve faster than structured proteins at the sequence level [37, 38]. This is reflected in the enrichment of lower mean interspecies amino acid conservation across all levels of constraint, even for the high CCRpct residues (Table 1). High percentiles of constraint ([95, 100]) were enriched with residues in disorder to order (D-to-O), disorder to disorder (D-to-D) and context dependent transitions upon binding with other protein partners, with higher OR values for those undergoing D-to-O and context dependent transitions. These observations align with what has been proposed in terms of the binding mechanisms for these structural transitions. D-to-O are defined by a single, well-defined, fully ordered binding configuration, mediated by a unique well-defined contact pattern that excludes ambiguities and is determined by the presence of binding motifs. Conversely, D-to-D transitions are defined by many different binding configurations, including alternative contact patterns, often with weak or redundant motifs. Context dependent transitions involve alternative binding configurations, which change with the cellular conditions and different partners [15]. Residues in regions driving LLPS present the highest association we observed, with OR 9.26 times more likely to be in the most highly constrained regions (percentiles [99, 100]) and with the longest average length of 75 amino acids (Figure 4). LLPS regions are generally mobile/disordered and involve long sequence stretches that orchestrate multiple and multivalent interactions with proteins and RNA for the formation of membrane-less organelles in the cell and, unsurprisingly, these two types of interactions are also observed with large odds of being highly constrained (ORs 3.16 and 4.82, respectively). Furthermore, 29 (53%) of the 54 proteins in our dataset containing annotated LLPS regions have amino acid sites with percentiles in [95, 100] in coincidence with such regions (Supplementary Table 5). When considering gnomAD2.1.1 per gene variant intolerance metrics, 34 out of these 54 (63%) LLPS driver proteins have pLI >= 0.9 (i.e. are ‘essential’ genes, extremely intolerant to pLoF variants in heterozygosity), while only 7 of the 54 are highly intolerant to missense changes (missense OEUF<=0.35) (Supplementary Table 5).
IV. ***Signalling regions.*** Residues in propeptides, signal peptides and transit signalling regions tend to be in regions more likely to have unconstrained to medium constrained percentiles in the interval [0, 60] (Figure 5 D). The very few highly constrained sites in pro-peptides and signal peptides show much lower conservation and shorter region length (Figure 4 and Table 1).
V. ***Post translationally modified positions (PTMs).*** The overall tendency is for sites that are post translationally modified to be more likely in constrained regions at lower percentiles (Figure 5 E). Given that these are specific sites with, in general, very short flanking motifs, it is expected that shorter regions, and therefore lower CCRpct, are associated with these positions. Also, some PTMs may not be functionally relevant and represent false positives [39]. Glycosylated sites tend to be more associated with being unconstrained or at low constraint (percentiles [0, 60]). The number of lipidated sites is very low, therefore statistically not significant for most of the CCRs percentile categories. However, they are less likely to be unconstrained (OR=0.62, 95% CI: 0.53-0.72). Other types of PTMs are more likely to be associated with highly constrained regions (percentiles [95, 99]). There are 482 highly constrained sites, 302 (62%) correspond to N-acetylations, and 79% are N-acetyl lysine. This is not surprising given the abundance of the latter modification. When comparing conservation and constraint for these sites, the behaviour is mixed. Phosphorylation sites tend to be less conserved in general for all constraint levels (Table 1), and a slightly higher prevalence is for sites with high constraint and low conservation. Lipidated sites tend to be more conserved and with low or high percentiles. Glycosylation and other PTMs are more associated with high conservation for all constraints and with slightly higher preference for the low CCRs high conservation combination.
VI. ***Other relevant sites and regions.*** The localisation of “other sites” and “other regions” was obtained from UniProt. This category corresponds to regions/positions of functional relevance for proteins, identified mostly from experimental evidence, that cannot be described by other feature annotations of UniProt. We also recorded coding DNA sequence (CDS) junctions, by translating the genomic coordinates of these sites obtained from Ensembl onto the corresponding amino acids in the UniProt sequences. As suspected, given their relevance, the protein sites in these three categories are more likely to be constrained at high percentiles (Figure 5 F) and also mostly related to high percentiles and high conservation (Table 1).
VII. ***Residues without annotations.*** 1.8M amino acids did not have any of the functional sites or regions or domains that we aggregated in the present study. These were very weakly associated with unconstrained and low constraint regions and particularly less associated with higher percentiles (Figure 5 G). They were slightly associated with all combinations of constraints and low conservation (Table 1), and slightly more with low CCRpct and low conservation. The 96.4K and 658.7K residues that are in CCRpct [50, 100] with low and high conservation, could be explained by functional features that still remain to be discovered and annotated for some proteins, or some domains that were difficult to delimit, creating fuzziness at their boundaries.

In summary, when bringing together the classifications regarding CCRpct and protein functional features, for the 9.8M positions that can be mapped to a CCRs percentile, it is possible to observe how the co-occurrence with certain functional features becomes more evident at higher percentiles (Figure 5 A vs B and Figure 4). The 23.6K most highly constrained residues in the human genome (CCRpct in [99, 100], 0.24% of the total mapped residues) correspond to, on average, the longest linear stretches depleted of tolerated variability, and strongly highlight positions involved in DNA/RNA binding, protein binding, catalytic sites and in driving LLPS.

### Amino acid sites with clinically interpreted variants and their CCR percentiles

We next investigated how residues with different types of clinically interpreted variants correlate with the different percentiles of constraint. For this purpose, we employed variants from the ClinVar database (https://www.ncbi.nlm.nih.gov/clinvar), and classified the amino acid positions in our dataset as pathogenic (including pathogenic/likely_pathogenic), benign (including benign/likely_benign) and/or VUS/conflicting (including variants of uncertain significance or with conflicting interpretations of pathogenicity), and followed a similar methodology for performing OR tests (Supplementary Figures 4 and 6) Pathogenic missense variants account for only 28,398 protein sites, with the majority of missense variants in ClinVar corresponding to VUS/conflicting interpretations, affecting 194,508 residues. Residue sites with pathogenic missense variants were observed as strongly associated with higher percentiles of constraint, in the intervals between [90, 100]. Surprisingly, these types of variants were also associated with unconstrained regions (percentile [0, 0]), although with lower odds. Sites with benign and VUS/conflicting missense variants had higher odds of being in unconstrained regions, although a few benign variants, affecting 476 amino acid sites, were also present in the top most highly constrained regions.

**Figure 6:**
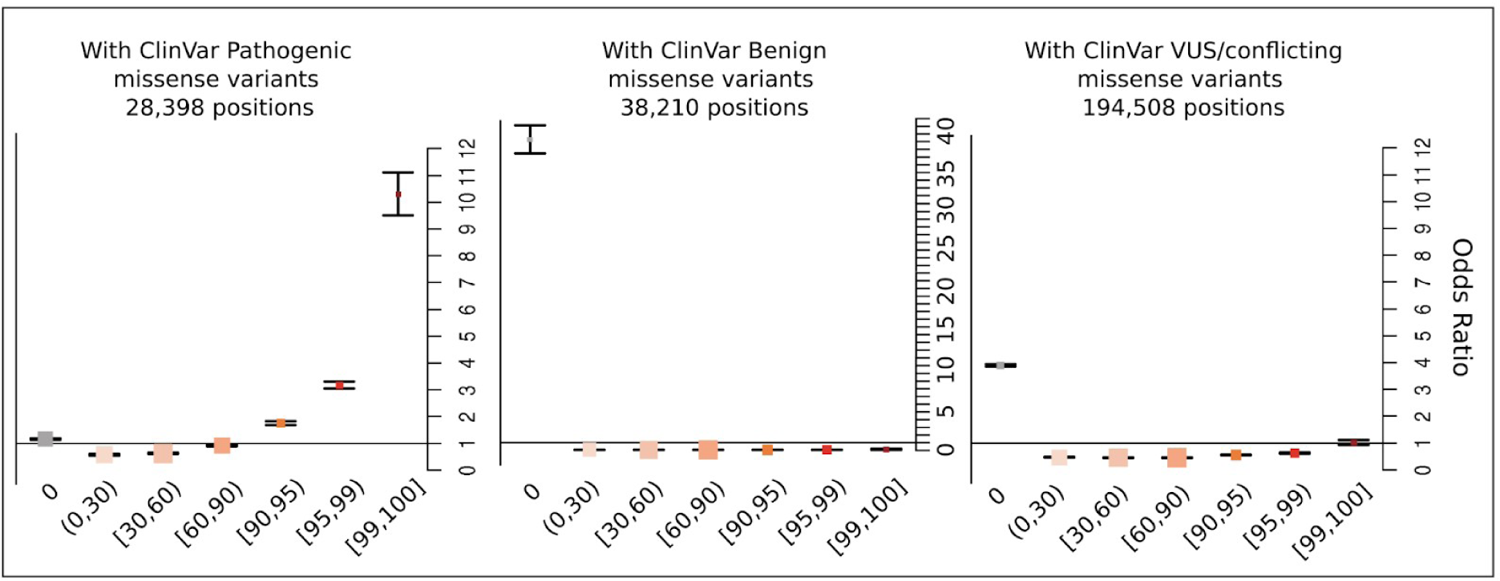
Propensity of co-occurrence of amino acid positions having clinically interpreted missense variants with the different CCRpct. The ORs with 95% CI based on two-tailed Fisher’s exact test, represent the odds that amino acid sites with a particular type of variant will co-occur in combination with one of the CCRs percentile categories, compared to the odds of having such a type of variant but with any of the other percentile categories. The vertical lines through the boxes give the length of the CI. The line at OR=1 is the line of no clear difference; boxes and intervals above this represent more likely co-occurrences, while boxes and intervals under the line represent the converse. See Supplementary Table 3 for all the p-values corresponding to the two and one tailed Fisher’s exact tests.

When comparing the distribution of residues with the three groups of variants with CCRpct and conservation scores, we observed that they are spread across all categories of conservation and CCRpct (Supplementary Table 4 and Supplementary Figure 3), but only the amino acid sites affected by pathogenic/likely_pathogenic were OR=1.63 times more likely to be more conserved and with high CCRpct.

Additionally, we assessed the co-occurrence of ClinVar variants with the 30 protein feature annotations (Supplementary Figures 6 and 7). Sites with missense pathogenic variants are mostly associated with being in domains, transmembrane regions, catalytic sites, metal binding, ligand binding, protein binding, disulphide bonds, DNA/RNA binding, linear motifs, LIPs, D-to-O, lipidation, other regions, and CDS junctions. Sites with missense benign variants were more likely to be disordered/mobile, low complexity, LLPS, signal, propeptide and transit peptides, and sites without any protein features. Sites with missense VUS/conflicting variants, were mostly associated with those regions generally more difficult to characterise: repeats, coiled-coils, LIPS, D-to-O and D-to-D transitions, phosphorylation sites, other regions of biological relevance annotated in UniProt, and sites without any feature.

**Figure 7:**
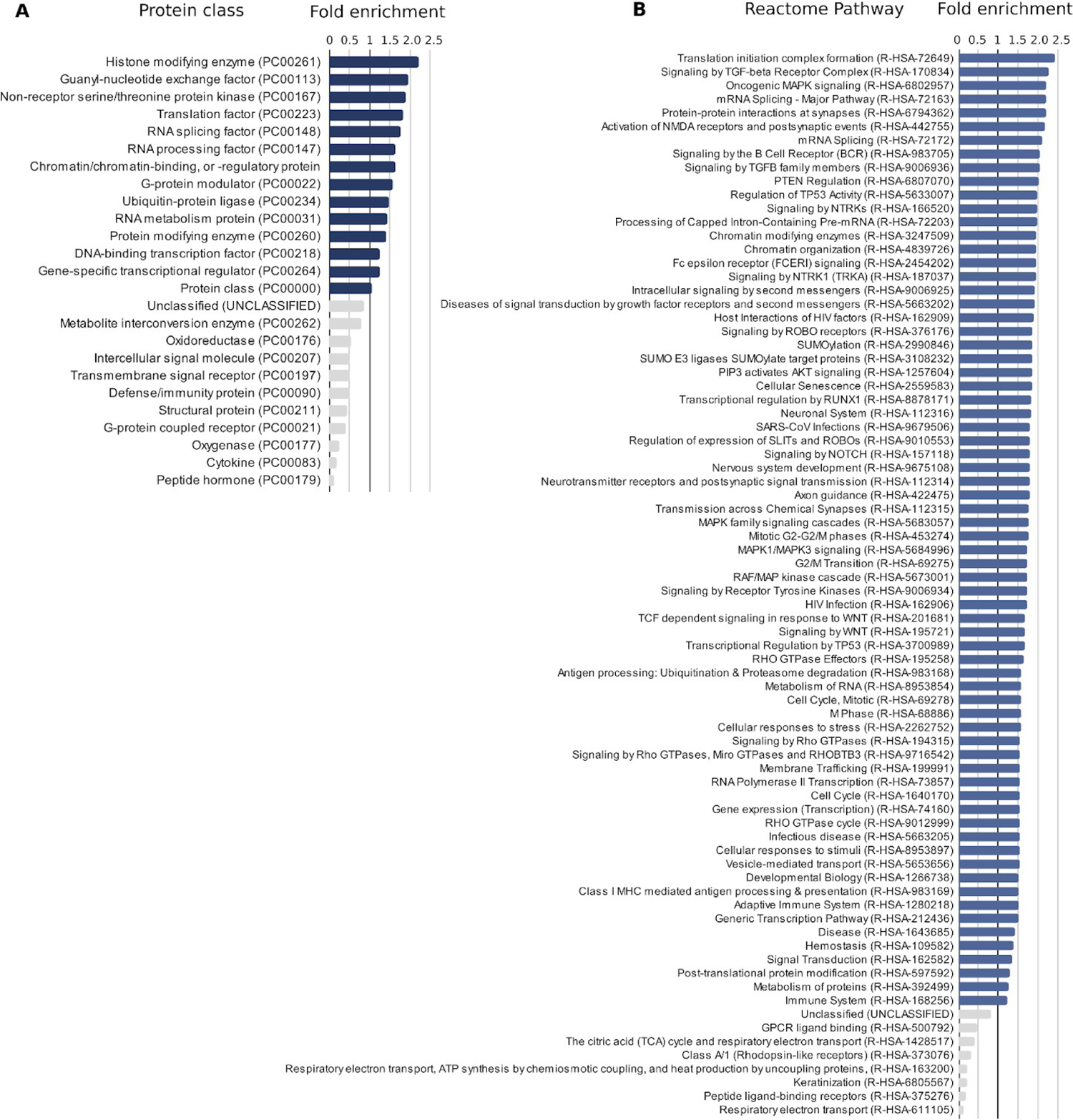
Gene Ontology enrichment test for A) protein class, and B) pathways annotated in Reactome, for proteins harbouring residues in highly constrained regions with percentiles in [95, 100]. The overrepresentation tests are based on multiple Fisher tests with Bonferroni correction, and only significant (p-value<0.05) terms are listed and ordered by fold of enrichment. Bars in blue correspond to over represented terms (>1 fold of enrichment).

LLPS driving regions presented a low number of sites with Pathogenic variants, while being highly associated with high CCRpct (see Figures 4 A and 5 E). This motivated us to further investigate the distribution of Pathogenic variants in the corresponding proteins, the biological processes where they are involved and the types of diseases they associate with. The Supplementary Table 5 presents this information, and Supplementary Table 6 summarises the list of clinical conditions and the amount of LLPS driving genes associated with them.

We observed the majority of LLPS driving genes (35 out of the 54, 63%) acting in different biological processes that facilitate DNA damage repair, epigenetic gene repression and RNA metabolism (transcription, splicing, polyadenylation, transport, translation, see column ‘Main biological process groups’ and the corresponding genes in Supplementary Table 5). The remaining 19 genes are involved in different processes including neuron cell growth, adhesion, axonogenesis and synaptogenesis, development, synaptic plasticity and regulation of neurotransmitter vesicles release; signal transduction pathways for cell survival, migration, proliferation, differentiation and apoptosis; protein degradation/recycling; cell cycle regulation; immune responses; nuclear transport; elasticity of organs and tissues; muscle structure and function and glomerular filtration in kidney.

34 (64%) of the 54 genes driving LLPS have pLI>=0.9 (i.e. high intolerance to loss of function in heterozygosity), in particular this is the case for genes involved in RNA metabolism. 43 of these LLPS proteins (79.6%) have amino acid positions which drive phase separation and are highly constrained for variation (CCRpct in [90, 100]).

24 LLPS genes (44% out of the total 54) have not yet been associated to any disease by protein changing variants by the time we consulted ClinVar. 14 of these 24 have pLI⩾0.9 and are associated to RNA metabolism (10 genes), DNA damage repair (1 gene), immune response (1 gene) and signal transduction for cell proliferation and differentiation (1 gene), all of them presenting multiple regions of high constraint (CCRpct [90, 100]).

We observed that 17 (31.5%) of the 54 genes were associated with severe early onset developmental disorders, including different organ malformations, oestrogen resistance with absence of sexual maturation or severe early onset immunodeficiency, with 10 of them in particular linked to neurodevelopmental disorders (Supplementary Table 5, column ‘Disease group’). 10 genes (18.5% of the 54) were associated with later onset diseases, mostly neurodegenerative but also affecting muscles and bone and triggering earlier menopause. 5 genes (9%) were associated with different cancers of pancreas, breast, uterus and prostate, lung and leukemia. There was also high incidence of associations with not yet described conditions (see Supplementary Table 6, ‘not provided’ or ‘not specified’).

### Overrepresentation of regions with high percentiles in GO protein classes and Reactome pathways

We investigated whether there was an enrichment of the types of proteins and biological pathways in the proteins having regions with high CCRpct. For this purpose, we submitted a ‘query list’ of 6,402 protein identifiers of sequences with percentiles in the interval [95, 100] to the PANTHER Classification System [40] to perform a Gene Ontology (GO) Overrepresentation Test for two categories of terms: ‘protein class’ and ‘Reactome pathways’. We used as the ‘reference list’ the 17,366 genes/proteins for which we have CCRs estimates; however, for only 17,022 PANTHER was able to assign GO terms.

Genes with [95, 100] CCRpct were enriched in 14 protein classes out of a total of 196 that were assigned to our lists of proteins. Genes with [95, 100] CCRpct were enriched in 70 Reactome pathways, out of a total of 2.482. Figure 9 A and B list the statistically relevant over- and under-represented protein classes and Reactome Pathways, respectively. Additionally, the 29 out of 54 (53%) LLPS proteins that have highly constrained amino acid sites in their LLPS driving regions are mainly involved in RNA metabolism, processing, splicing, transcription and translation initiation (Panther Protein Class in Supplementary Table 5).

### Exploring some examples: U2AF2, the splicing factor U2AF 65 kDa subunit, and SLC12A2, the solute carrier family 12 member 2

Aggregating different protein annotations such as inter-species conservation, human variability and constraint, functional features, 3D structure and presence of clinically interpreted variants can help understand why variants have different propensities in different contexts. We have chosen two examples of proteins for illustration, the first of which illustrates a protein with highly disordered/mobile regions which have nevertheless been constrained during human evolution, although the conservation across species is patchy. The second example is a transmembrane protein in which the functional ion channel residues are highlighted as highly constrained by the CCRs and also a disordered region of 20 amino acids which are variable across species but depleted of tolerated variants and include a cluster of pathogenic variants.

The first example is *U2AF2*, the 65 kDa subunit of the U2 auxiliary splicing factor U2AF (Figure 7), a protein that is highly constrained against variability. This is an essential splicing factor that recognizes the polypyrimidine-tract (Py) 3’ splice-site signal in pre-mRNA and initiates spliceosome assembly in the nucleus [41]. *U2AF2* is highly constrained for missense and LoF variability in gnomAD2.1 (missense OEUF=0.31, missense Z-score=4.21, pLI=1, LOEUF=0.133), suggesting essentiality for humans. There is partial structural data for this protein, derived from 3 separate PDB files, each containing a different part of the protein. The protein contains a low complexity arginine-serine rich motif (RS) at positions 27–62, which has been proposed to initiate liquid-liquid phase separation (LLPS) to form nuclear speckle drops in the nucleus, bringing together pre-mRNAs and the proteins of the spliceosome [42]. Further along the sequence, the region 85-112 is a UHM ligand motif (ULM) that has been shown responsible for the interactions with U2AF1 (also known as U2AF35) the 35kDa subunit of the splicing factor *U2AF* [43]. The two central RNA recognition motifs (RRM) are shown in the Pfam domains panel and central dashed box of Figure 7, with protein structures on top. These regions bind to the Py-tract signal in the pre-RNA, are highly mobile (light grey shading regions in the Pfam domains panel) [44, 45], and are connected by a flexible/disordered linker region (231–258) that modulates the binding specificity for the proper Py-tracts in pre-mRNA [46]. The third RRM domain, known as U2AF Homology Motif, UHM, is atypical and has lost its RNA-binding ability, but interacts with splicing factor 1 (SF1) (right dashed box with protein structures on top) [46, 47]. The protein has long, moderate to highly constrained regions (CCRs panels) that co-localize with the U2AF2-U2AF1 binding interface, the three RRMs, the flexible linker and also with regions involved in LLPS and/or linear interacting peptides (pink and violet rectangles in the “Other protein features” panel). A few variants have been reported in the literature as associated with a developmental disorder and different types of cancer (represented with lollipops above Pfam domains in Figure 7) and only a VUS is reported in ClinVar. All the pathological missense variants represent drastic physiochemical changes, and affect highly constrained and highly conserved sites.

*U2AF2* illustrates how CCRpct and conservation both highlight that this protein is not only essential, but also has functional protein sites along its length. The clinical data supports this hypothesis, with variants associated with developmental disorders and cancers.

The second example is shown in Figure 8, which illustrates how CCR data, combined with protein function information, can highlight regions with potential functions that have yet to be determined. The gene is *SLC12A2*, which encodes for the solute carrier family 12 member 2 protein, a Na+, K+ and 2Cl− cotransporter 1 (NKCC1), which plays a critical role in the homeostasis of K+ enriched endolymph in the membranous labyrinth of the inner ear. NKCC1 subunits are ion channels that work as homodimers, and each protein monomer comprises a transmembrane domain (TMD), cytosolic N- and C-terminal domains (NTD and CTDs, respectively) and extracellular flexible loops stabilised by disulphide bonds. The dimeric interface involves interactions between TMDs and CTDs [50, 51].

**Figure 8:**
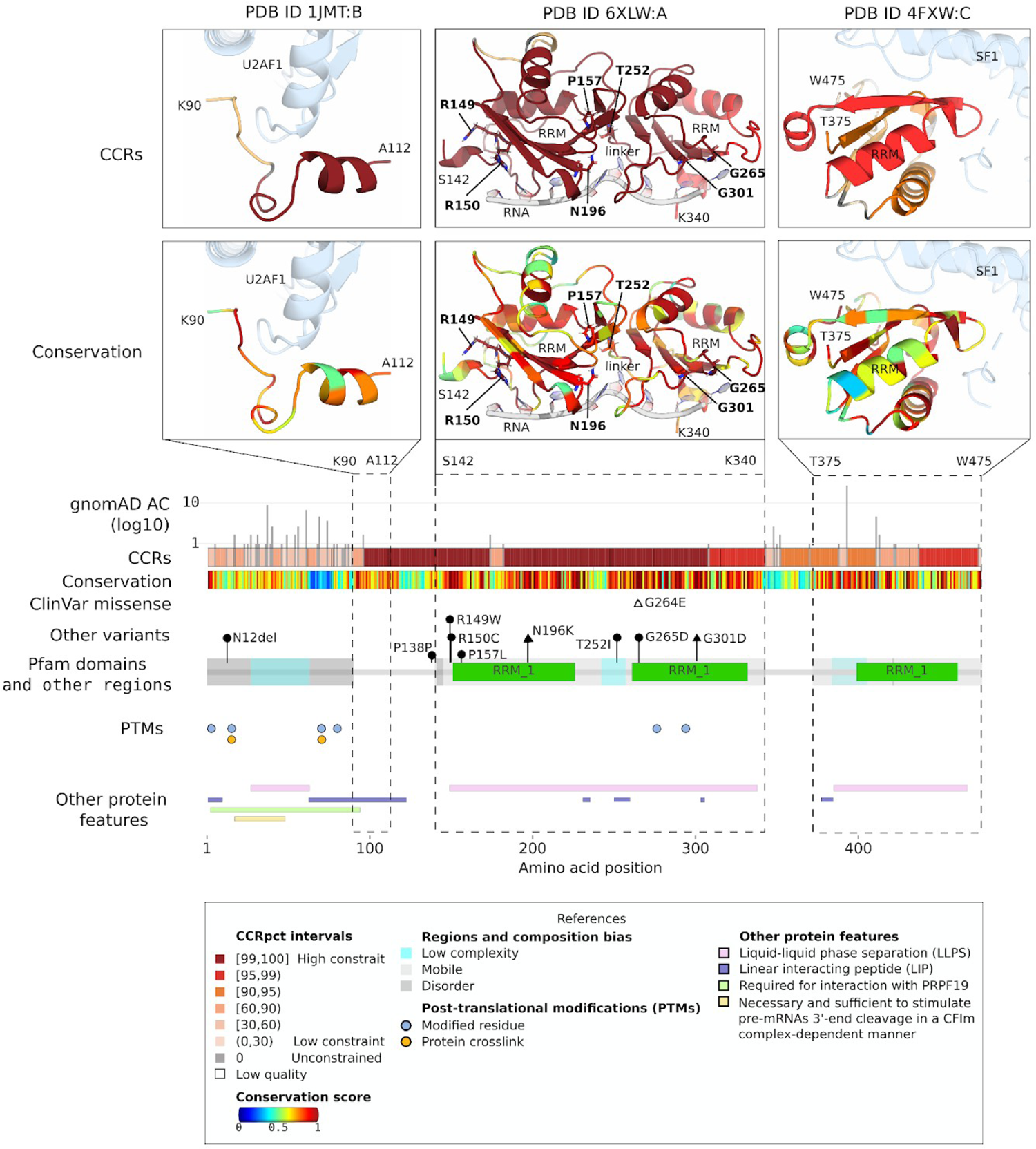
An example of a protein with long regions highly constrained for variability: *U2AF2,* the 65 kDa subunit of the U2 auxiliary splicing factor U2AF (also known as *U2AF65*, UniProtKB: P26368, Ensembl: ENST00000308924). From middle to bottom the different panels represent, by amino acid positions, gnomAD3.0 allele counts (AC), CCRpct, species conservation score, Pfam domain with disorder/mobility and low-complexity regions, post-translationally modified sites (PTMs), other protein features, as listed in the Figure. Pfam domains correspond to: ‘RRM_1’=RNA recognition motif, RNP-1. On top of these panels, the dashed rectangles call out regions of the protein with available PDB structures. The lollipops above the Pfam domains depict positions with *de novo* variants reported in bibliography as associated to developmental disorders (N12del, P138P, R149W, R150C, P157L, T252I and G265G with black circles) [48], acute myeloid leukaemia (N196K), colon adenocarcinoma and castration-resistant prostate carcinoma (G301D) (black triangles) [49]. The unfilled triangle represents a variant of uncertain significance reported in ClinVar (G264E). By-residue ScoreCons conservation scores were obtained from VarSite. Low-complexity, mobility, disorder LLPS, and LIP annotations were obtained from the MobiDB database. PTMs and other interacting regions were obtained from UniProtKB. Interacting proteins in the PDB structures are shown in pale blue, with their interacting side chain shown in stick representation.

*SLC12A2* is overall highly constrained for missense and LoF variability in gnomAD2.1 (missense OEUF=0.78, missense Z-score=2.4, pLI=0.96, LOEUF=0.31), suggesting essentiality for humans. The protein structures (Figure 8), reveal that the highest CCRpct in this protein (in the range [90,99)) corresponds to functionally important regions: residues in pore-lining helices (involved in the ion flow through the channel), the dimer interface, and also a ‘not so obviously relevant’ disordered and lowly conserved (scores < 0.4) region in the C-terminal domain. Intriguingly, this last region, comprising amino acids 977 to 993, is fully encoded by exon 21 of *SLC12A2* and coincides with the boundaries of a CCR ranked with a moderate percentile of 94 (small dashed rectangle in Figure 8). It has previously been noted that this region, whose functionality still remains unclear, is unique to SLC12A2 and is not shared with the other proteins in the SLC12 family [52], suggesting that it might confer a specific functional characteristic to this protein.

Seven pathogenic missense variants have been reported in ClinVar for this protein. Two of them, associated with Delpire-McNeill neurodevelopmental syndrome, are in the TMD in highly conserved and low-medium constrained residues: N376I lining the pore (conservation=1, CCRs pct=37.68) and A327V adjacent to a pore-lining helix (conservation=0.72, CCRs pct=74.95). The other five: E979K, E980K, D981Y, P988T and P988S cause deafness and sensorineural hearing loss, and cluster in the moderately constrained region encoded by exon 21. Furthermore, functional assays in cultured cells showed that applying the variants E979K, D981Y and P988T, or skipping exon 21, significantly decreases chloride influx mediated by the SLC12A2 protein [53]. All this evidence and the moderately high CCRpct for this disordered and lowly conserved region, highlight its putative relevance for the function of this protein. Both the U2AF2 and SLC12A2 proteins described above also serve to exemplify different scenarios for amino acid sites where high CCRs percentiles go hand in hand with high conservation, and the converse where high CCRs percentiles go with low conservation, and vice versa.

## Discussion

In the present work we extended the characterisation of constrained coding regions in the human genome, by accurately fine-mapping these regions and their level of constraint from the Human Build 38 genomic coordinates to protein sequence coordinates in 17,366 human UniProt canonical sequences, totalling about 9.8 million amino acid positions. Furthermore, aggregating protein functional annotations, available for these positions, allowed us to analyse the distribution and correlation of the different levels of constraint and inter-species conservation with different protein features.

For the catalytic sites and interactions with different partners (small molecules, proteins, DNA/RNA, metals), we observed the expected associations between high percentiles of constraint and high conservation scores. This is in concordance with the observations of Havrilla et al. [1] that domains enriched with the most highly constrained regions were involved in ion transport and in different DNA/RNA interactions (like zinc fingers, helicases and translation factors). Additionally, we observed that the unconstrained (i.e. with gnomAD3.0 variants) or lowly constrained (i.e. average shorter regions depleted of variants) regions were mostly associated with signalling regions (signal, propeptides and transit peptides), low complexity, glycosylation sites, and with more mixed inter-species conservation scores.

Surprisingly, the transmembrane regions showed little if any enrichment for highly constrained regions, but slightly higher enrichment for medium constraint (CCRpct [60,90)), i.e. on average shorter regions. Perhaps, this reflects the lipid environment where variants between the hydrophobic amino acids are common.

Among the unexpected results, we observed that disulphide bond cysteines were more prone to lie within regions with low to medium percentiles of constraint (CCRpct in (0,90)). Disulphide bonds are covalent tertiary interactions important for stabilising protein folds and/or performing physiologically relevant redox activity and hence highly conserved in evolution [54]. We hypothesise that the association with lower-medium CCRpct (i.e. average shorter regions depleted of variants, with mean length=20 amino acidos) reflects the fact that the formation of such bonds requires only the presence of short motifs involving only the cysteines and their immediate flanking residues [55]).

Perhaps the most unexpected results we observed were related to disordered and mobile regions in proteins, showing dual enrichment for unconstrained/lowly constrained and also for highly constrained percentiles, mostly in sites with low conservation, and this might relate to the multiplicity of functions, or “flavours”, of disorder that such regions can present, which depend on their length, composition and location in proteins [14, 56]. Disordered proteins and regions are able to fulfil a variety of tasks: they can serve as flexible linkers between structured regions or flexible binding sites for ligands, they can undergo disorder-order transitions upon binding to other proteins through specific molecular recognition features (MoRFs) within longer disordered regions, they can also have short linear motifs that work as targets for post-translational modifications or cell signalling, or longer regions which promote molecular recognition and protein-protein interactions. The characterisation of the dynamic of IDPs/IDRs has led to the identification of their plausible role in regulating enzymatic activity [57] and has also been useful to investigate ligand selection for developing drugs [58]. This motivates the necessity of characterising disordered proteins and regions, for discovering the function and relevant mechanism where they are involved.

In the present work, in particular, residues involved in D-to-O, context dependent transitions and in driving LLPS showed association with high constraints, for conserved and also unconserved sites. Furthermore, our results suggest that LLPS amino acids are strongly associated with constrained regions ranked with the highest percentiles ([95, 100]) and that such regions are, on average, the longest stretches (75 amino acids in length) depleted of protein changing variants across the human coding genome. However, it was remarkable that amino acid positions in regions driving LLPS were not significantly associated with Pathogenic protein altering variants, considering that previous works have observed these genes frequently related to cancer, autism spectrum disorders, neurodegeneration, and infectious diseases [21–23]. Here, apart from such associations, we also observed 64% of the LLPS driving genes were highly constrained for loss-of-function in heterozygosity and 31.5% were involved in diseases with profound impact in the normal human postnatal and early development, with a high prevalence of neurodevelopmental disorders. Also, 44% of the 54 genes were not associated with any disease and 14 of them, mostly associated to RNA metabolism, were highly constrained for loss-of-function variation. We hypothesise that such genes could possibly affect embryonic viability. In this sense, it is worth to mention that LLPS drives the formation of membrane-less organelles important for organising and regulating key cellular processes such as transcription, splicing, translation, chromosome condensation, synapsis and downstream signalling [59–62], all essential for tightly regulating the differential expression of genes, ensuring cell survival, correct differentiation into different tissues, and for the development and function of the neuronal and immune systems.

The scientific community is beginning to untangle the complexity of interactions and regulations involved in LLPS, and evidence shows that these protein condensates do not follow classical rules of molecular recognition [63]. Furthermore, the current tools that attempt to predict the clinical relevance of a specific sequence variant have been developed mostly based on the characteristics of folded protein regions [64] making it difficult to understand the effect of variants affecting intrinsically disordered/mobile and liquid-liquid phase separating regions.

Most of the significantly enriched GO terms for proteins with highly constrained regions (CCRpct in [95, 100]) were related to RNA-processing, DNA binding, protein-protein interactions, and enzymatic activities. This is in concordance with our observations that amino acid sites functionally annotated as binding DNA/RNA and/or proteins, in catalytic sites and in LIPs, and/or driving LLPS, are among the ones with the greatest odds of being highly constrained, i.e. in the, on average, longest regions in the human genome intolerant to variability.

Our results also complement and extend what the authors of the CCRs model [1] have derived before by analysing the co-occurrence with Pfam domains and observing that the highly constrained regions are involved in ion transport and in different DNA/RNA interactions (like zinc fingers, helicases and translation factors), but also that about 30% of these highly constrained regions did not correspond to any protein Pfam [3] domain.

Undoubtedly, the sequencing of genetically more diverse human populations will refine some CCRs further, but the data presented here has significant clinical utility. The key challenge for clinical genomics is interpreting the pathogenicity of rare variants. Identifying whether a rare variant lies within a defined constrained region of the protein facilitates consequence interpretation especially for novel variants absent from existing genomic databases.

We emphasise that combining interspecies and intraspecies (human population) conservation can help to highlight regions of individual genes that have appeared more recently in evolution or confer some degree of uniqueness/specificity to an individual paralogue. This data has the potential to facilitate the discovery of new associations between genes/variants with previously unknown phenotypes. CCRs also highlight many highly constrained regions currently not linked to any Mendelian disease. This may indicate mutations in these regions are lethal to humans or are sufficiently rare that they have not yet been identified [1].

Furthermore, many protein functional sites still remain to be characterised and currently lack sufficient functional annotations, and also the particularly difficult cases, where flexible linkers/disordered regions are poorly characterised and/or can be poorly conserved across species while being constrained, at different levels, in human populations. Our mapping of CCRs to amino acids helps to define these regions in proteins more accurately and could contribute to the further annotation of these challenging regions.

## Methods

### Generation of the CCRs based on gnomAD3

To obtain the CCRs we ran the pipeline developed by [1] (https://quinlan-lab.github.io/ccr/examples/updates) but employing the dataset of gnomAD [26] version 3.0 and the corresponding files for the coordinates of the human genome in version GRCh38 [65] (https://www.ncbi.nlm.nih.gov/grc). We used the Variant Effect Predictor (VEP) [66] of Ensembl [25] version 101. We also followed the recommendations of the authors to only consider genetic variants from autosomes and chromosome X, and avoid those in conflicting genomic regions - i.e. where there are segmental duplications and/or high identity with other genomic regions (>=90% identity) or with low sequencing coverage. In the same line of recommendations, we ran the weighting of the regions for autosomes and X chromosome separately, but merged both output files into one for performing the mapping of the coordinates of the regions to protein amino acids (see *Mapping the CCRs to protein amino acids*).

### Mapping the CCRs to protein amino acids

We developed an in-house pipeline in R that uses the ‘*ensembldb*’ Bioconductor R package [67] to map the genomic coordinates of CCRs boundaries, and all the coding bases in between, to the Ensembl v101 transcripts which are part of the GENCODE [68] basic set version 35. This was to ensure we were including complete and well annotated relevant transcripts. For those amino acid sites where the corresponding codon had constrained and unconstrained bases, we assigned such amino acid sites as unconstrained. We then obtained the sequence identifiers that crosslink Ensembl transcripts and proteins and UniProtKB proteins [24] by querying the APIs (application programming interfaces) of both databases. Finally, the CCRpct were accurately transferred to the amino acids in UniProtKB sequences by downloading the corresponding protein sequences from Ensembl and UniProtKB and performing Blastp local alignments [69] requesting 100% sequence identity (perfect match). The workflow is summarised in Figure 10 A, the corresponding scripts of the pipeline are available in this repository https://github.com/marciaah/CCRStoAAC.

**Figure 9:**
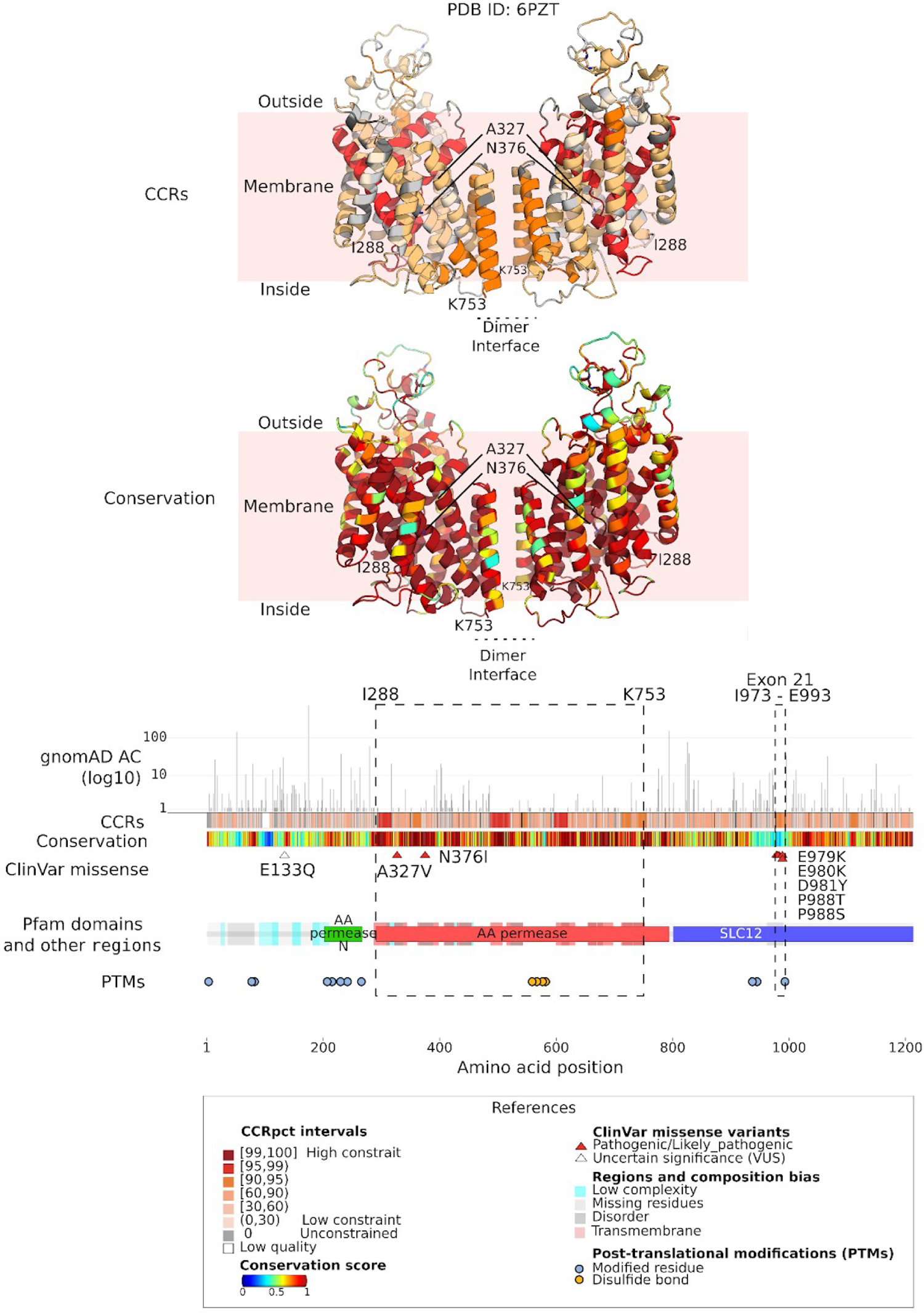
An example where CCRs highlight regions important for protein function: the pore lining helices and dimer interface in the transmembrane region of *SLC12A2* (NKCC1) solute carrier family 12 member 2 (UniProtKB: P55011-1, Ensembl: ENST00000262461) and a peculiar disordered region of this protein encoded by its exon 21 and with a cluster of pathogenic/likely_pathogenic variants related to deafness and hearing loss [53]. From middle to bottom the plots in the horizontal panels represent, by amino acid position, gnomAD3.0 allele counts (AC), CCRpct, species conservation scores, sites with ClinVar missense variants, Pfam domains with disorder/mobility, low-complexity and transmembrane regions and post-translationally modified sites (PTMs). Pfam domains correspond to: ‘AA permease N’=Amino acid permease N-terminal, ‘AA permease’= Amino acid permease, ‘SLC12’=Solute carrier family 12. Over these domains, the dashed rectangles call out the transmembrane region of the protein characterised in the 6PZT PDB structure as a homodimer. This structure is coloured by CCRpct and by conservation and displayed on the top panels. The location in the structure of the pathogenic/likely_pathogenic ClinVar variants A327V and N376I is depicted with triangles and squares, respectively. The small dashed rectangle fully encloses exon 21 (amino acids 977-993). Residue conservation scores, as calculated by ScoreCons, were obtained from VarSite, domains from Pfam, low-complexity, mobility and disorder from MobiDB, transmembrane regions and PTMs from UniProtKB.

**Figure 10:**
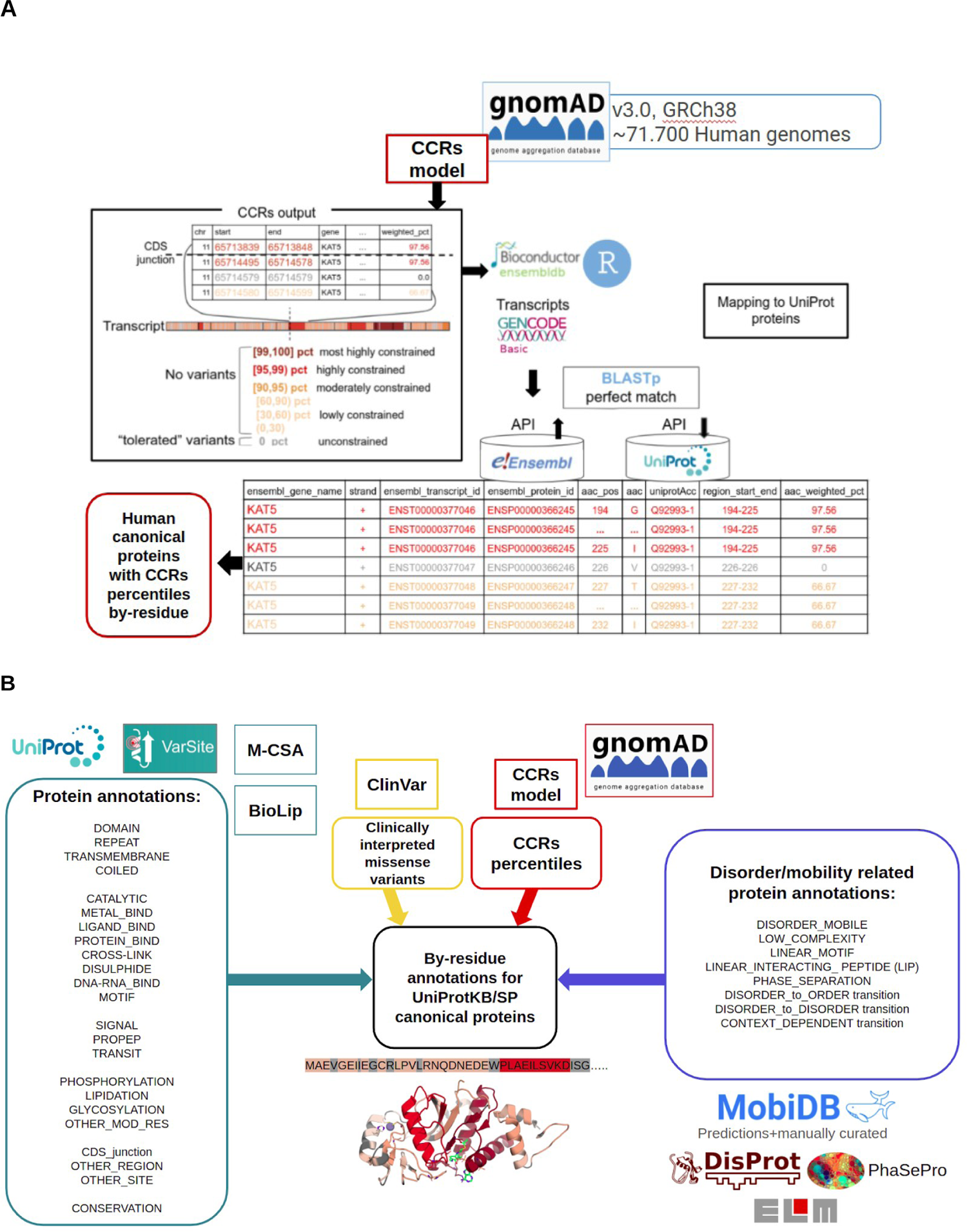
A) Flowchart showing the different databases and tools employed for mapping the CCRs in genomic coordinates to the amino acid coordinates in UniProtKB protein sequences. B) Flowchart presenting the different resources and databases employed for aggregating 30 general protein feature annotations and conservation score (blue-green boxes), clinically interpreted variants (yellow boxes), CCRpct (red boxes) and disorder/mobile related protein feature annotations (blue boxes). MOTIF and LINEAR_MOTIF were combined in a single category. UniProtKB/SP= UniProtKB Swiss Prot

### Aggregation of protein feature annotations and clinically interpreted variants

We developed our own pipeline in R for fetching different protein features and functional annotations from multiple resources. For this purpose, we captured annotations and based our analysis only on UniProtKB/SwissProt canonical sequences because such sequences are the main references for annotations in the databases we employed. An overview of the databases and features that we included are presented in Figure 10 B, and obtained as described more in detail in Supplementary Methods.

### Gene Ontology enrichment tests

The GO statistical overrepresentation tests were performed using the PANTHER classification system [40] (http://www.pantherdb.org/tools/index.jsp, PANTHER version 17.0 release 22-02-2022, with Reactome version 65), submitting the list of genes of interest (e.g. those presenting regions with percentiles in [95, 100]) and using as “reference list” only those genes for which we were able to map CCRpct.

### Odds ratios tests for enrichment

We performed four different Odds ratio (OR) test analyses to measure the enrichment of amino acid sites presenting different combinations of CCRpct, conservation, protein features and ClinVar variants:

I. *CCRpct and presence of each one of the 30 protein features:* we binary assigned whether or not an amino acid site had any of the protein features (see Figure 11 for full list) and a CCRpct in any of the 7 bins: unconstrained=[0]; low-medium constraint= (0,30), [30,60) and [60,90); moderately constrained= [90,95), highly constrained= [95,99) and most highly constrained= [99, 100].
II. *CCRpct and conservation with the presence of protein features:* we binary classified the amino acid sites as having or not any of the 30 protein features and any of the 6 combinations: a) CCRs unconstrained (0 pct) and conservation score ≤0.5, b) CCRs unconstrained (0 pct) and conservation score >0.5, c) CCRs in (0,50] pct and conservation score ≤0.5, d) CCRs in (0,50] pct and conservation score >0.5, e) CCRs in (50,100] pct and conservation score ≤0.5, f) CCRs in (50,100] pct and conservation score >0.5.
III. *CCRpct and presence of ClinVar variants:* amino acid sites were binary assigned whether or not they had “pathogenic/likely_pathogenic”, “benign/likely_benign” or “VUS/conflicting interpretations of pathogenicity” variants and a CCRpct in any of the 7 bins mentioned in (I).
IV. *CCRpct and conservation with the presence of ClinVar variants:* we classified residues according to whether they had or not any of the 3 groups of variants as described in (III) and any of the 6 combinations of CCRpct and conservation as described in (II).

For the four enrichment analyses, contingency tables were constructed counting amino acid sites with the different classifications (See Supplementary Methods: *OR tests for enrichment* for further details) and the OR were calculated using two-tailed and one-tailed Fisher’s exact tests [70] for obtaining the corresponding P-values and 95% confidence intervals (CI 95%). It is worth clarifying that when counting residues we did not request exclusivity in the intersections, i.e. a residue with a given CCRpct can intersect with being in DOMAIN, DISORDER_MOBILE and DNA-RNA_BIND and hence will contribute to the cells in the three corresponding contingency tables.

## Acknowledgments

This work was funded by the Cambridge NIHR Biomedical Research Centre (BRC-1215-200014) and the 3D-FunSites Project (Wellcome Trust Ref. number 221327/Z/20/Z). The Wellcome Trust, the European Bioinformatics Institute (EMBL-EBI) and the Medical Research Council have also funded research infrastructure. We thank Prof. Raymond’s and Prof. Thornton’s group members for their valuable comments and suggestions.

## Data Availability Statement

The code and data used for the present analysis is provided in GitHub repositories, as mentioned in the Methods section: https://github.com/marciaah/CCRStoAAC and https://github.com/marciaah/CCRStoAAC-output. The authors welcome requests for additional information regarding the material presented in this paper.

