## Supplementary Figures for "Mapping the Constrained Coding Regions in the human genome to their corresponding proteins"

**Supplementary figures 1 to 7**

**Title:** *Mapping the Constrained Coding Regions in the human genome to their corresponding proteins*

Marcia A. Hasenahuer[1,2,5], Alba Sanchis-Juan[3,4,6], Roman A. Laskowski[1], James A. Baker[1], James D. Stephenson[1], Christine A. Orengo[5], F. Lucy Raymond[2,4], Janet M. Thornton[1]

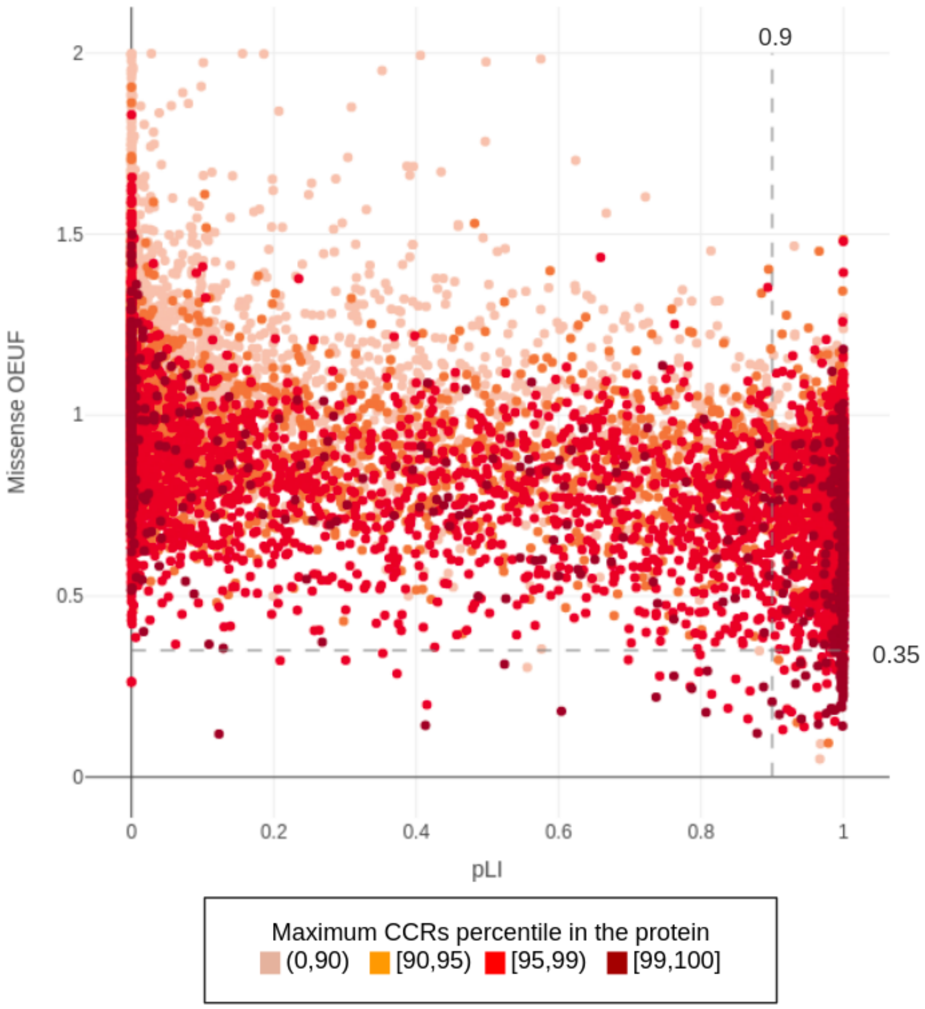


**Supplementary figure 1: Comparison of pLI and missense OEUF and the maximum CCRs percentile by protein.** Thresholds for missense OEUF (≤0.35) and pLI (≥0.9) scores, showing the proteins (dots) with the highest constraints for missense and pLoF respectively, are depicted with dashed lines. The lower right corner, enclosed by both thresholds, groups genes highly constrained for pLoF and missense variants

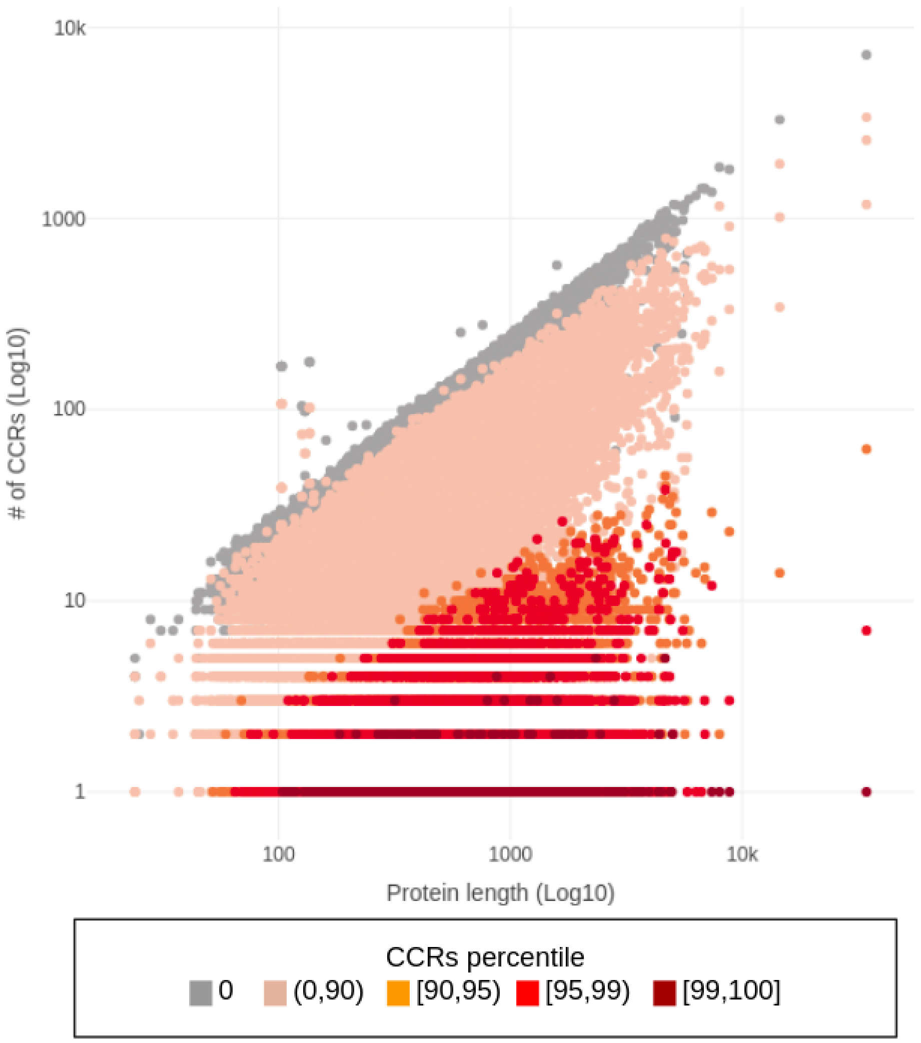


**Supplementary Figure 2:** Comparison of full length of proteins and the number of regions in each CCRs percentile that they have.

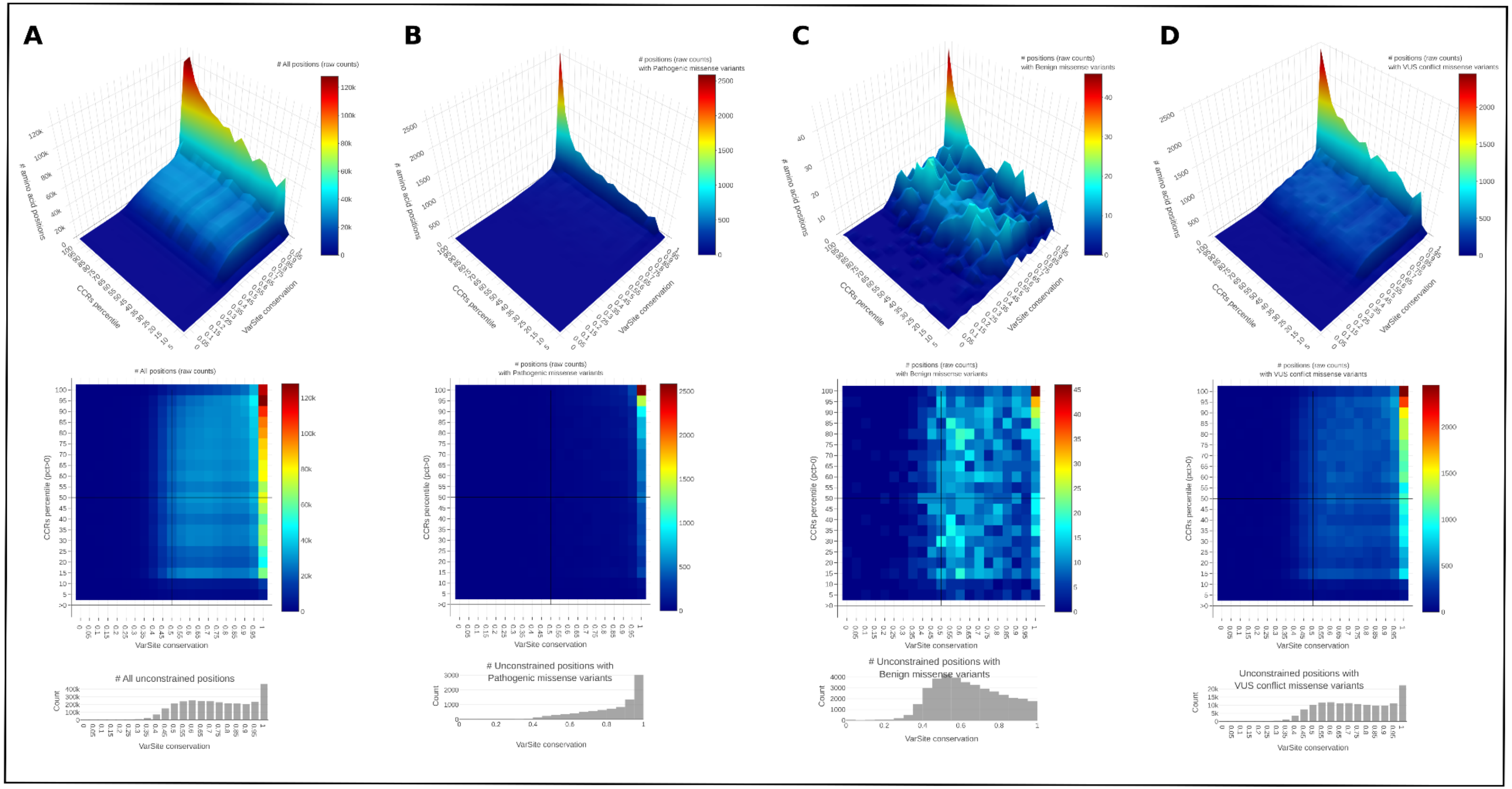

A
B
C
D
### All positions (raw counts)
### positions (raw counts) with Pathogenic missense variants
### positions (raw counts) with Benign missense variants
### positions (raw counts) with VUS conflict missense variants
### amino acid positions
CCRs percentile
VarSite conservation
CCRs percentile (pct>0)
### All unconstrained positions
### Unconstrained positions with Pathogenic missense variants
### Unconstrained positions with Benign missense variants
Unconstrained positions with VUS conflict missense variants
Count

**Supplementary Figure 3: Distribution of the raw count of amino acid positions grouped by different categories of CCRs percentiles and conservation scores**. A) all positions, and those affected by B) Pathogenic/Likely_pathogenic, C) Benign/Likely_benign and D) VUS/conflictive variants, in DOMAIN

| CCRs pct | [T][T] | [T][F] | [F][T] | [F][F] | OR | CI 95% | Fisher p-value |
|---|---|---|---|---|---|---|---|
| 0,0] | 1065616 | 2618209 | 2003314 | 4139901 | 0.84 | 0.84-0.84 | 0 * |
| 0,30) | 483897 | 3199928 | 915102 | 5227648 | 0.86 | 0.86-0.87 | 0 * |
| 30,60) | 803415 | 2880410 | 1414811 | 4728123 | 0.93 | 0.93-0.94 | 0 * |
| 60,90) | 961599 | 2722226 | 1450553 | 4692355 | 1.14 | 1.14-1.15 | 0 * |
| 90,95) | 198797 | 3485028 | 226160 | 5916047 | 1.49 | 1.48-1.5 | 0 * |
| 95,99) | 157074 | 3526751 | 125600 | 6016533 | 2.13 | 2.12-2.15 | 0 * |
| 99,100] | 15221 | 3668604 | 8386 | 6133688 | 3.03 | 2.95-3.12 | 0 * |

in TRANSMEMBRANE

| CCRs pct | [T][T] | [T][F] | [F][T] | [F][F] | OR | CI 95% | Fisher p-value |
|---|---|---|---|---|---|---|---|
| [0,0] | 116477 | 272003 | 2952453 | 6486863 | 0.94 | 0.93-0.95 | 8.0e-66 * |
| (0,30) | 52412 | 336068 | 1346586 | 8091985 | 0.94 | 0.93-0.95 | 2.1e-42 * |
| [30,60) | 88398 | 300082 | 2129825 | 7309086 | 1.01 | 1-1.02 | 5.4e-03 * |
| [60,90) | 100808 | 287672 | 2311342 | 7127536 | 1.08 | 1.07-1.09 | 1.6e-94 * |
| [90,95) | 17748 | 370732 | 407209 | 9030502 | 1.06 | 1.05-1.08 | 4.0e-14 * |
| [95,99) | 11883 | 376597 | 270791 | 9166717 | 1.07 | 1.05-1.09 | 6.8e-12 * |
| [99,100] | 881 | 387599 | 22726 | 9414687 | 0.94 | 0.88-1.01 | 8.2e-02 |

in DISORDER-FLEXIBLE

| CCRs pct | [T][T] | [T][F] | [F][T] | [F][F] | OR | CI 95% | Fisher p-value |
|---|---|---|---|---|---|---|---|
| [0,0] | 619721 | 1192921 | 2449209 | 5565293 | 1.18 | 1.18-1.18 | 0.0e+00 * |
| (0,30) | 281918 | 1530724 | 1117080 | 6896904 | 1.14 | 1.13-1.14 | 0.0e+00 * |
| [30,60) | 405809 | 1406833 | 1812414 | 6201835 | 0.99 | 0.98-0.99 | 3.8e-11 * |
| [60,90) | 395676 | 1416966 | 2016474 | 5997809 | 0.83 | 0.83-0.83 | 0.0e+00 * |
| [90,95) | 62563 | 1750079 | 362394 | 7651042 | 0.75 | 0.75-0.76 | 0.0e+00 * |
| [95,99) | 43232 | 1769410 | 239442 | 7773865 | 0.79 | 0.79-0.8 | 0.0e+00 * |
| [99,100] | 5051 | 1807591 | 18556 | 7994695 | 1.2 | 1.17-1.24 | 1.8e-30 * |

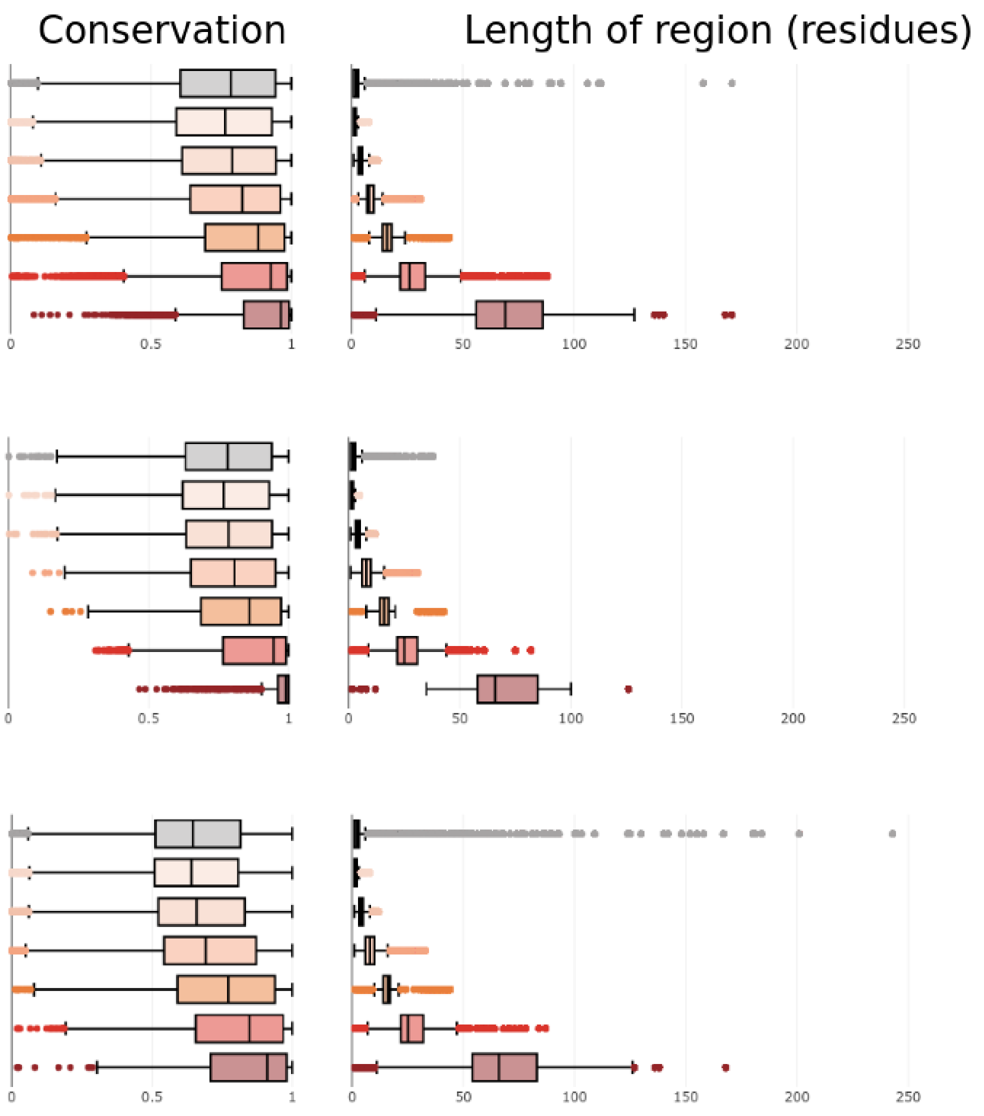

**in METAL BINDING**

| CCRs pct | [T][T] | [T][F] | [F][T] | [F][F] | OR | CI 95% | Fisher p-value |
|---|---|---|---|---|---|---|---|
| [0,0] | 151735 | 366320 | 2917195 | 6392489 | 0.91 | 0.9-0.91 | 2.8e-212 * |
| (0,30) | 64822 | 453233 | 1334177 | 7974774 | 0.85 | 0.85-0.86 | 3.8e-301 * |
| [30,60) | 108517 | 409538 | 2109707 | 7199546 | 0.9 | 0.9-0.91 | 1.0e-184 * |
| [60,90) | 134660 | 383395 | 2277490 | 7031762 | 1.08 | 1.08-1.09 | 5.0e-135 * |
| [90,95) | 30406 | 487649 | 394551 | 8913523 | 1.41 | 1.39-1.43 | 0.0e+00 * |
| [95,99) | 25465 | 492590 | 257209 | 9050728 | 1.82 | 1.8-1.84 | 0.0e+00 * |
| [99,100] | 2777 | 515278 | 20830 | 9287010 | 2.4 | 2.31-2.5 | 0.0e+00 * |

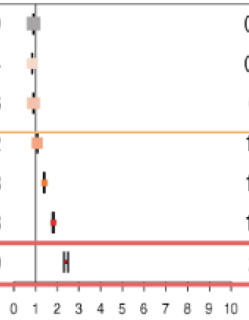

**in LIGAND BINDING**

| CCRs pct | [T][T] | [T][F] | [F][T] | [F][F] | OR | CI 95% | Fisher p-value |
|---|---|---|---|---|---|---|---|
| [0,0] | 170839 | 421294 | 2898091 | 6337611 | 0.89 | 0.88-0.89 | 0.0e+00 * |
| (0,30) | 75395 | 516738 | 1323603 | 7911343 | 0.87 | 0.87-0.88 | 5.5e-263 * |
| [30,60) | 127159 | 464974 | 2091065 | 7144193 | 0.93 | 0.93-0.94 | 2.1e-97 * |
| [60,90) | 152403 | 439730 | 2259747 | 6975484 | 1.07 | 1.06-1.08 | 3.3e-106 * |
| [90,95) | 33870 | 558263 | 391087 | 8842954 | 1.37 | 1.36-1.39 | 0.0e+00 * |
| [95,99) | 29248 | 562885 | 253426 | 8980432 | 1.84 | 1.82-1.86 | 0.0e+00 * |
| [99,100] | 3334 | 588799 | 20273 | 9213488 | 2.57 | 2.48-2.67 | 0.0e+00 * |

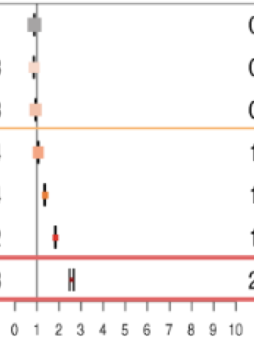

**in CATALYTIC SITES**

| CCRs pct | [T][T] | [T][F] | [F][T] | [F][F] | OR | CI 95% | Fisher p-value |
|---|---|---|---|---|---|---|---|
| [0,0] | 779 | 2697 | 3068151 | 6756257 | 0.64 | 0.59-0.69 | 7.1e-31 * |
| (0,30) | 448 | 3028 | 1398550 | 8425077 | 0.89 | 0.81-0.98 | 2.2e-02 * |
| [30,60) | 726 | 2750 | 2217497 | 7606468 | 0.91 | 0.83-0.98 | 1.8e-02 * |
| [60,90) | 1018 | 2458 | 2411132 | 7412806 | 1.27 | 1.18-1.37 | 1.9e-10 * |
| [90,95) | 263 | 3213 | 424694 | 9398023 | 1.81 | 1.59-2.05 | 1.4e-17 * |
| [95,99) | 213 | 3263 | 282461 | 9540057 | 2.2 | 1.91-2.53 | 1.3e-23 * |
| [99,100] | 29 | 3447 | 23578 | 9798839 | 3.5 | 2.34-5.03 | 1.9e-08 * |

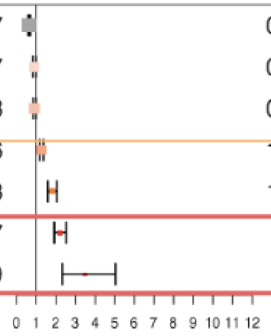

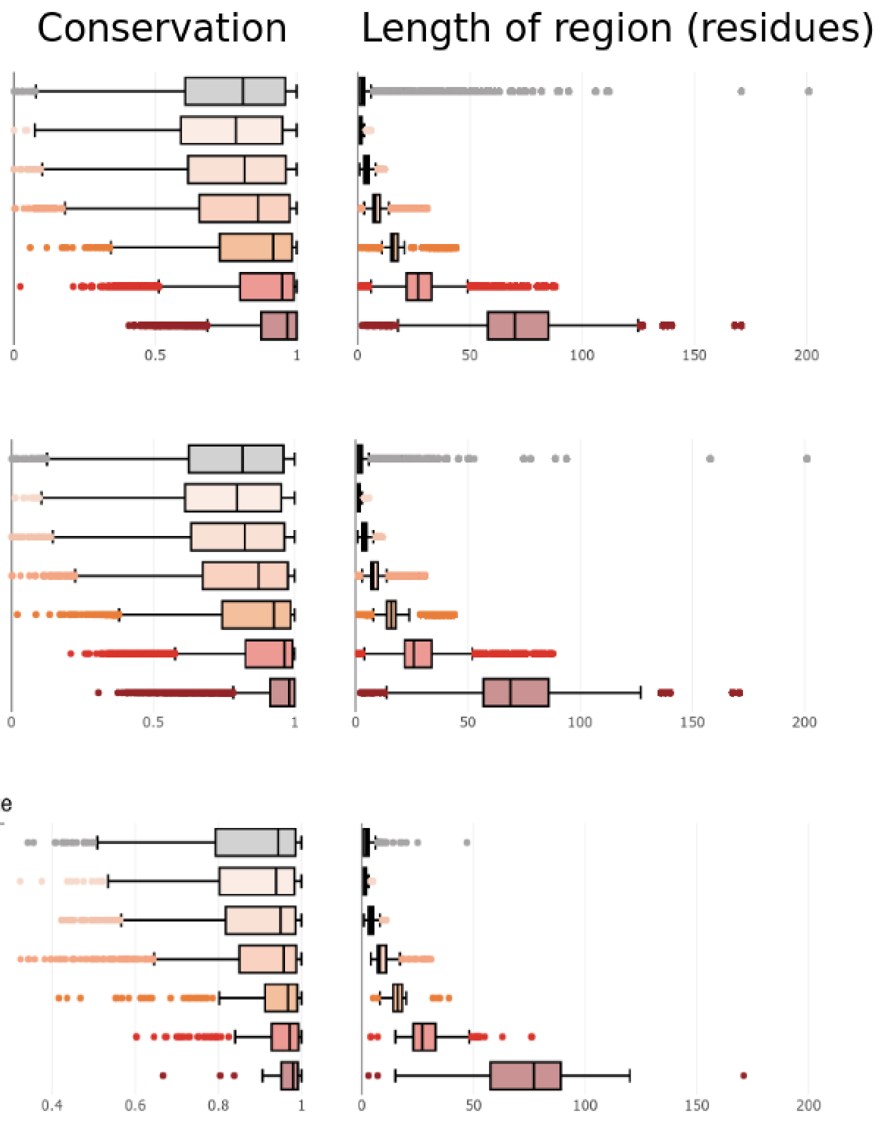

DISORDER-to-ORDER transition

| CCRs pct | [T][T] | [T][F] | [F][T] | [F][F] | OR | CI 95% | Fisher p-value |
|---|---|---|---|---|---|---|---|
| [0,0] | 16258 | 43037 | 3052672 | 6715810 | 0.83 | 0.82-0.85 | 1.2e-91 * |
| (0,30) | 7218 | 52077 | 1391780 | 8375950 | 0.83 | 0.81-0.86 | 4.4e-49 * |
| [30,60) | 11626 | 47669 | 2206597 | 7561492 | 0.84 | 0.82-0.85 | 2.4e-69 * |
| [60,90) | 16222 | 43073 | 2395928 | 7372123 | 1.16 | 1.14-1.18 | 4.9e-56 * |
| [90,95) | 3699 | 55596 | 421258 | 9345633 | 1.48 | 1.43-1.53 | 1.4e-103 * |
| [95,99) | 3980 | 55315 | 278694 | 9487994 | 2.45 | 2.37-2.53 | 0.0e+00 * |
| [99,100] | 485 | 58810 | 23122 | 9743476 | 3.48 | 3.17-3.8 | 4.0e-113 * |

DISORDER-to-DISORDER transition

| CCRs pct | [T][T] | [T][F] | [F][T] | [F][F] | OR | CI 95% | Fisher p-value |
|---|---|---|---|---|---|---|---|
| [0,0] | 59277 | 128995 | 3009653 | 6629854 | 1.01 | 1-1.02 | 1.5e-02 * |
| (0,30) | 26981 | 161291 | 1372017 | 8266736 | 1.01 | 0.99-1.02 | 2.4e-01 |
| [30,60) | 40965 | 147307 | 2177258 | 7461866 | 0.95 | 0.94-0.96 | 1.1e-17 * |
| [60,90) | 45171 | 143101 | 2366979 | 7272090 | 0.97 | 0.96-0.98 | 1.7e-08 * |
| [90,95) | 8376 | 179896 | 416581 | 9221331 | 1.03 | 1.01-1.05 | 7.7e-03 * |
| [95,99) | 6850 | 181422 | 275824 | 9361885 | 1.28 | 1.25-1.31 | 3.4e-82 * |
| [99,100] | 822 | 187450 | 22785 | 9614836 | 1.85 | 1.72-1.98 | 3.1e-56 * |

CONTEXT DEPENDENT transition

| CCRs pct | [T][T] | [T][F] | [F][T] | [F][F] | OR | CI 95% | Fisher p-value |
|---|---|---|---|---|---|---|---|
| [0,0] | 238 | 621 | 3068692 | 6758333 | 0.84 | 0.72-0.98 | 2.7e-02 * |
| (0,30) | 88 | 771 | 1398910 | 8427334 | 0.69 | 0.55-0.86 | 6.1e-04 * |
| [30,60) | 217 | 642 | 2218006 | 7608576 | 1.16 | 0.99-1.35 | 6.1e-02 |
| [60,90) | 246 | 613 | 2411904 | 7414651 | 1.23 | 1.06-1.43 | 6.2e-03 * |
| [90,95) | 16 | 843 | 424941 | 9400393 | 0.42 | 0.24-0.69 | 1.1e-04 * |
| [95,99) | 46 | 813 | 282628 | 9542507 | 1.91 | 1.39-2.57 | 8.6e-05 * |
| [99,100] | 8 | 851 | 23599 | 9801435 | 3.9 | 1.68-7.73 | 1.3e-03 * |

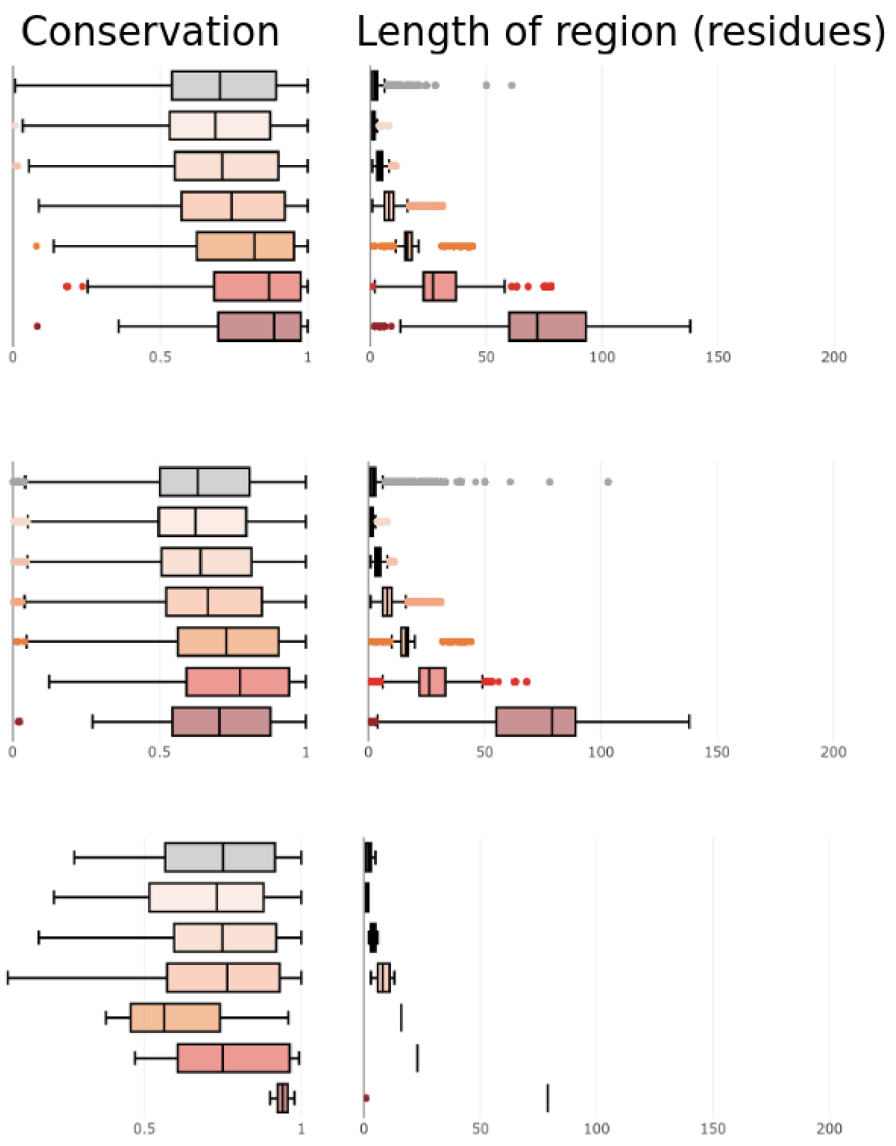

**Supplementary figure 4: Comparison of OR tests, conservation and length of regions by protein feature.** For each protein feature or functional site, the plots show: the contingency tables with the sites counts, OR, 95% CI and p-values along with boxplots of distribution of CCRs regions length and interspecies amino acid conservation. Stars (*) highlight statistically significant ORs (p-value ≤ 0.05) PLACEHOLDER FIGURES, all the plots by protein feature will go here.

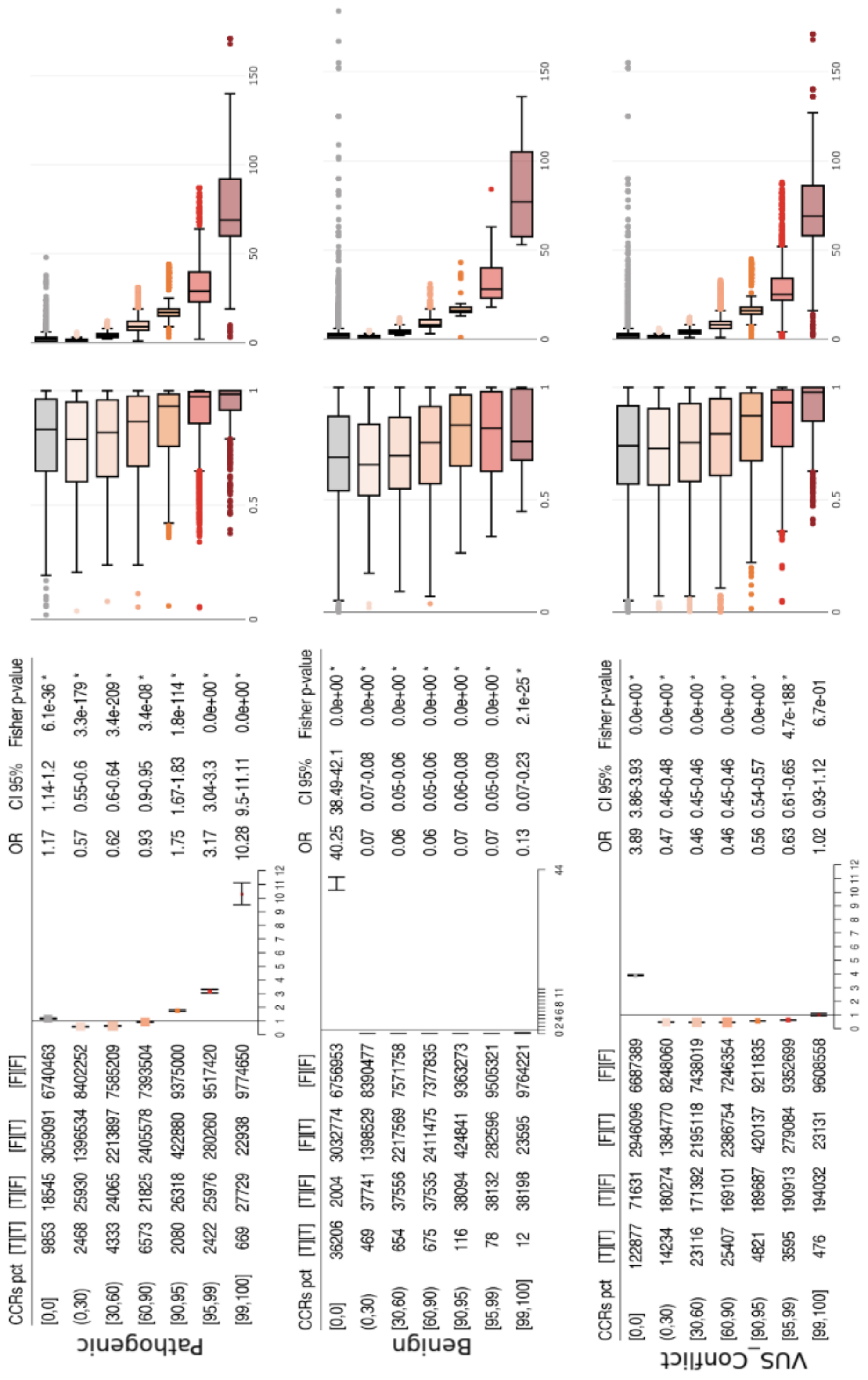

**Supplementary figure 5: Comparison of OR tests, conservation and length of regions by group of ClinVar variants**. For each category of ClinVar missense, the plots show contingency tables with the sites counts, OR, 95% CI and p-values along with boxplots of distribution of CCRs regions length and interspecies amino acid conservation. Stars (*) highlight statistically significant ORs (p-value ≤ 0.05)

**Odds Ratios (OR) forest plot**
**DOMAIN residues and ClinVar variants**

| Clinvar type | [T][T] | [T][F] | [F][T] | [F][F] | OR | CI 95% | Fisher p-value |
|---|---|---|---|---|---|---|---|
| P_miss | 23584 | 128314 | 4814 | 96686 | 3.69 | 3.57-3.81 | 0.0e+00 |
| B_miss | 18501 | 133397 | 19709 | 81792 | 0.58 | 0.56-0.59 | 0.0e+00 |
| VC_miss | 115106 | 36792 | 79402 | 22101 | 0.87 | 0.85-0.89 | 1.5e-46 |
| P_syn | 294 | 88308 | 137 | 58935 | 1.43 | 1.17-1.77 | 4.6e-04 |
| B_syn | 70776 | 17826 | 48251 | 10822 | 0.89 | 0.87-0.91 | 9.1e-18 |
| VC_syn | 18329 | 70273 | 11216 | 47857 | 1.11 | 1.08-1.14 | 1.1e-15 |
| P_sgsl | 13323 | 1895 | 9316 | 1547 | 1.17 | 1.09-1.26 | 2.7e-05 |
| B_sgsl | 164 | 15054 | 165 | 10698 | 0.71 | 0.56-0.88 | 1.9e-03 |
| VC_sgsl | 1760 | 13458 | 1430 | 9433 | 0.86 | 0.8-0.93 | 1.1e-04 |
| P_other | 20288 | 5076 | 14907 | 5168 | 1.39 | 1.33-1.45 | 1.8e-47 |
| B_other | 337 | 25027 | 896 | 19179 | 0.29 | 0.25-0.33 | 1.4e-93 |
| VC_other | 5096 | 20268 | 4673 | 15402 | 0.83 | 0.79-0.87 | 2.6e-16 |

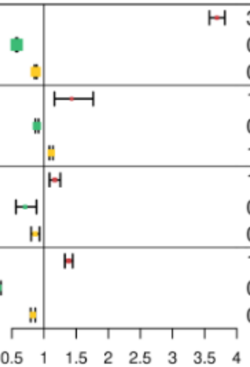


**Odds Ratios (OR) forest plot**
**TRANSMEMBRANE residues and ClinVar variants**

| Clinvar type | [T][T] | [T][F] | [F][T] | [F][F] | OR | CI 95% | Fisher p-value |
|---|---|---|---|---|---|---|---|
| P_miss | 2137 | 6554 | 26261 | 218458 | 2.71 | 2.58-2.85 | 5.9e-279 |
| B_miss | 885 | 7806 | 37325 | 207386 | 0.63 | 0.59-0.68 | 3.3e-42 |
| VC_miss | 6019 | 2672 | 188489 | 56228 | 0.67 | 0.64-0.7 | 7.2e-60 |
| P_syn | 15 | 5543 | 416 | 141701 | 0.92 | 0.51-1.54 | 9.0e-01 * |
| B_syn | 4331 | 1227 | 114696 | 27423 | 0.84 | 0.79-0.9 | 4.0e-07 |
| VC_syn | 1258 | 4300 | 28287 | 113831 | 1.18 | 1.1-1.26 | 8.5e-07 |
| P_sgsl | 513 | 79 | 22126 | 3363 | 0.99 | 0.78-1.27 | 9.0e-01 * |
| B_sgsl | 12 | 580 | 317 | 25172 | 1.64 | 0.84-2.93 | 9.3e-02 * |
| VC_sgsl | 68 | 524 | 3122 | 22367 | 0.93 | 0.71-1.2 | 6.1e-01 * |
| P_other | 1053 | 333 | 34142 | 9911 | 0.92 | 0.81-1.04 | 1.8e-01 * |
| B_other | 32 | 1354 | 1201 | 42852 | 0.84 | 0.57-1.2 | 4.0e-01 * |
| VC_other | 322 | 1064 | 9447 | 34606 | 1.11 | 0.97-1.26 | 1.1e-01 * |

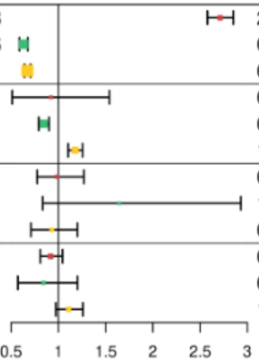


**Odds Ratios (OR) forest plot**
**REPEAT residues and ClinVar variants**

| Clinvar type | [T][T] | [T][F] | [F][T] | [F][F] | OR | CI 95% | Fisher p-value |
|---|---|---|---|---|---|---|---|
| P_miss | 1825 | 15941 | 26573 | 209072 | 0.9 | 0.86-0.95 | 3.8e-05 |
| B_miss | 2250 | 15516 | 35960 | 199677 | 0.81 | 0.77-0.84 | 1.8e-21 |
| VC_miss | 14320 | 3446 | 180188 | 55452 | 1.28 | 1.23-1.33 | 1.2e-37 |
| P_syn | 29 | 10080 | 402 | 137164 | 0.98 | 0.65-1.43 | 1.0e+00 * |
| B_syn | 7769 | 2340 | 111258 | 26314 | 0.79 | 0.75-0.82 | 4.3e-22 |
| VC_syn | 2405 | 7704 | 27140 | 110431 | 1.27 | 1.21-1.33 | 4.6e-22 |
| P_sgsl | 1485 | 198 | 21154 | 3244 | 1.15 | 0.99-1.35 | 7.4e-02 * |
| B_sgsl | 12 | 1671 | 317 | 24081 | 0.55 | 0.28-0.97 | 4.1e-02 |
| VC_sgsl | 188 | 1495 | 3002 | 21396 | 0.9 | 0.76-1.05 | 1.8e-01 * |
| P_other | 2016 | 493 | 33179 | 9755 | 1.2 | 1.09-1.33 | 3.0e-04 |
| B_other | 60 | 2449 | 1173 | 41757 | 0.87 | 0.66-1.13 | 3.4e-01 * |
| VC_other | 472 | 2037 | 9297 | 33637 | 0.84 | 0.75-0.93 | 6.7e-04 |

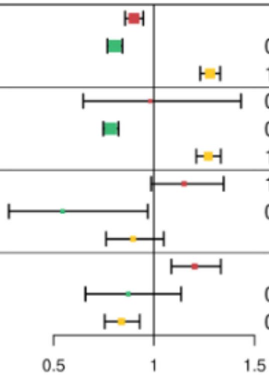


**Odds Ratios (OR) forest plot**
**DISORDER_FLEXIBLE residues and ClinVar variants**

| Clinvar type | [T][T] | [T][F] | [F][T] | [F][F] | OR | CI 95% | Fisher p-value |
|---|---|---|---|---|---|---|---|
| P_miss | 4971 | 47656 | 23427 | 177362 | 0.79 | 0.76-0.82 | 1.7e-48 |
| B_miss | 9375 | 43252 | 28835 | 171941 | 1.29 | 1.26-1.33 | 1.4e-83 |
| VC_miss | 40041 | 12586 | 154467 | 46315 | 0.95 | 0.93-0.98 | 4.4e-05 |
| P_syn | 98 | 30389 | 333 | 116855 | 1.13 | 0.89-1.42 | 2.8e-01 * |
| B_syn | 25221 | 5266 | 93806 | 23387 | 1.19 | 1.16-1.23 | 1.5e-26 |
| VC_syn | 5475 | 25012 | 24070 | 93122 | 0.85 | 0.82-0.87 | 4.1e-24 |
| P_sgsl | 4514 | 732 | 18125 | 2710 | 0.92 | 0.84-1.01 | 7.1e-02 * |
| B_sgsl | 82 | 5164 | 247 | 20588 | 1.32 | 1.02-1.71 | 3.2e-02 |
| VC_sgsl | 670 | 4576 | 2520 | 18315 | 1.06 | 0.97-1.17 | 1.9e-01 * |
| P_other | 8430 | 3001 | 26765 | 7247 | 0.76 | 0.72-0.8 | 3.1e-27 |
| B_other | 591 | 10840 | 642 | 33366 | 2.83 | 2.52-3.18 | 1.9e-68 |
| VC_other | 2709 | 8722 | 7060 | 26952 | 1.19 | 1.13-1.25 | 5.1e-11 |

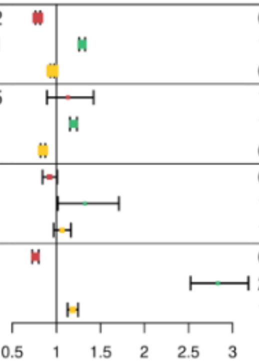

**Supplementary figure 6: Contingency tables that were built for performing the odds ratios (OR) tests, comparing each protein feature co-occurrence with different types of clinically interpreted variants from ClinVar**.

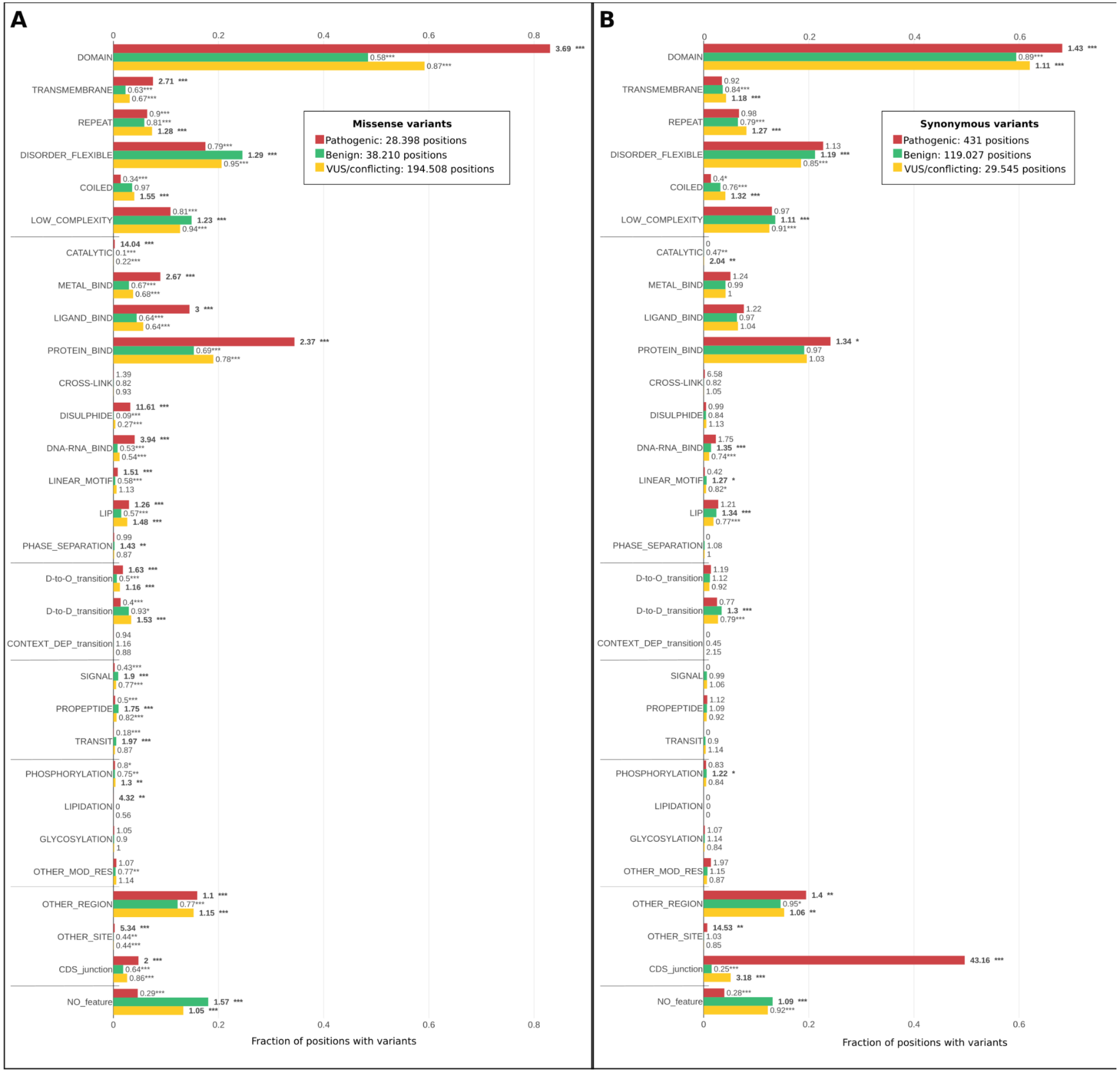
A
Missense variants
Pathogenic: 28.398 positions
Benign: 38.210 positions
VUS/conflicting: 194.508 positions
B
Synonymous variants
Pathogenic: 431 positions
Benign: 119.027 positions
VUS/conflicting: 29.545 positions
DOMAIN
TRANSMEMBRANE
REPEAT
DISORDER_FLEXIBLE
COILED
LOW_COMPLEXITY
CATALYTIC
METAL_BIND
LIGAND_BIND
PROTEIN_BIND
CROSS-LINK
DISULPHIDE
DNA-RNA_BIND
LINEAR_MOTIF
LIP
PHASE_SEPARATION
D-to-O_transition
D-to-D_transition
CONTEXT_DEP_transition
SIGNAL
PROPEPTIDE
TRANSIT
PHOSPHORYLATION
LIPIDATION
GLYCOSYLATION
OTHER_MOD_RES
OTHER_REGION
OTHER_SITE
CDS_junction
NO_feature
Fraction of positions with variants

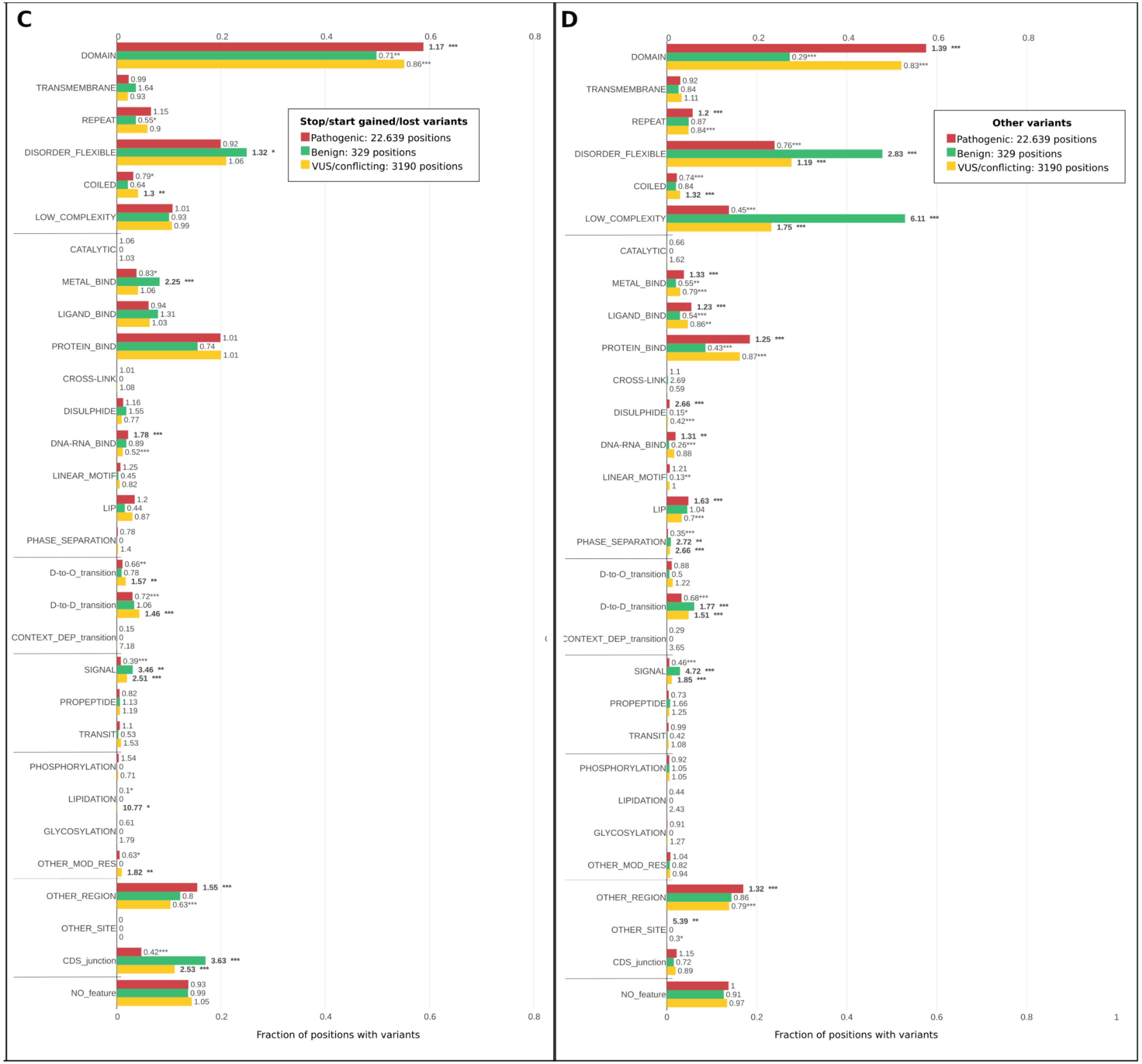
C
Stop/start gained/lost variants
Pathogenic: 22.639 positions
Benign: 329 positions
VUS/conflicting: 3190 positions
DOMAIN 1.17 *** 0.71** 0.86***
TRANSMEMBRANE 0.99 1.64 0.93
REPEAT 1.15 0.55* 0.9
DISORDER_FLEXIBLE 0.92 1.32 * 1.06
COILED 0.79* 0.64 1.3 **
LOW_COMPLEXITY 1.01 0.93 0.99
CATALYTIC 1.06 0 1.03
METAL_BIND 0.83* 2.25 *** 1.06
LIGAND_BIND 0.94 1.31 1.03
PROTEIN_BIND 1.01 0.74 1.01
CROSS-LINK 1.01 0 1.08
DISULPHIDE 1.16 1.55 0.77
DNA-RNA_BIND 1.78 *** 0.89 0.52***
LINEAR_MOTIF 1.25 0.45 0.82
LIP 1.2 0.44 0.87
PHASE_SEPARATION 0.78 0 1.4
D-to-O_transition 0.66** 0.78 1.57 **
D-to-D_transition 0.72*** 1.06 1.46 ***
CONTEXT_DEP_transition 0.15 0 7.18
SIGNAL 0.39*** 3.46 ** 2.51 ***
PROPEPTIDE 0.82 1.13 1.19
TRANSIT 1.1 0.53 1.53
PHOSPHORYLATION 1.54 0 0.71
LIPIDATION 0.1* 0 10.77 *
GLYCOSYLATION 0.61 0 1.79
OTHER_MOD_RES 0.63* 0 1.82 **
OTHER_REGION 1.55 *** 0.8 0.63***
OTHER_SITE 0 0 0
CDS_junction 0.42*** 3.63 *** 2.53 ***
NO_feature 0.93 0.99 1.05
0 0.2 0.4 0.6 0.8
Fraction of positions with variants
D
Other variants
Pathogenic: 22.639 positions
Benign: 329 positions
VUS/conflicting: 3190 positions
DOMAIN 1.39 *** 0.29*** 0.83***
TRANSMEMBRANE 0.92 0.84 1.11
REPEAT 1.2 *** 0.87 0.84***
DISORDER_FLEXIBLE 0.76*** 2.83 *** 1.19 ***
COILED 0.74*** 0.84 1.32 ***
LOW_COMPLEXITY 0.45*** 6.11 *** 1.75 ***
CATALYTIC 0.66 0 1.62
METAL_BIND 1.33 *** 0.55** 0.79***
LIGAND_BIND 1.23 *** 0.54*** 0.86**
PROTEIN_BIND 1.25 *** 0.43*** 0.87***
CROSS-LINK 1.1 2.69 0.59
DISULPHIDE 2.66 *** 0.15* 0.42***
DNA-RNA_BIND 1.31 ** 0.26*** 0.88
LINEAR_MOTIF 1.21 0.13** 1
LIP 1.63 *** 1.04 0.7***
PHASE_SEPARATION 0.35*** 2.72 ** 2.66 ***
D-to-O_transition 0.88 0.5 1.22
D-to-D_transition 0.68*** 1.77 *** 1.51 ***
CONTEXT_DEP_transition 0.29 0 3.65
SIGNAL 0.46*** 4.72 *** 1.85 ***
PROPEPTIDE 0.73 1.66 1.25
TRANSIT 0.99 0.42 1.08
PHOSPHORYLATION 0.92 1.05 1.05
LIPIDATION 0.44 0 2.43
GLYCOSYLATION 0.91 0 1.27
OTHER_MOD_RES 1.04 0.82 0.94
OTHER_REGION 1.32 *** 0.86 0.79***
OTHER_SITE 5.39 ** 0 0.3*
CDS_junction 1.15 0.72 0.89
NO_feature 1 0.91 0.97
0 0.2 0.4 0.6 0.8 1
Fraction of positions with variants

**Supplementary Figure 7: Amino acid positions affected with clinically interpreted variants assessed by their location in the different protein features or functional sites**. Each bar represents the fraction from the total positions with a feature that present variants assigned in ClinVar as having any of three types: Pathogenic/likely_pathogenic, Benign/Likely_benign or variant of uncertain significance or with conflicting interpretations of pathogenicity (VUS/conflicting ), for four groups of variants: A) missense, B) synonymous, C) stop/start gained/lost and D) other variants.
The numbers next to the bars are the OR, calculated with a 95% CI, and represent the odds that protein positions in a particular feature/functional site will co-occur in combination with one of the three types of missense variants, compared to the odds of having such feature/functional site but with any of the other two types of missense of variants (See Supplementary Figure 6 for further information on how the contingency tables were constructed). Stars denote the statistical significance (p-value) according to a two-tailed Fisher's exact test, the more stars the higher the significance (no star, not significant: p-value > 0.05; *: 0.05 ≥ p-value > 0.01; **: 0.01 ≥ p-value > 0.001; ***: p-value ≤ 0.001). Cases where OR>1 and 95% CI>1 with p-value ≤ 0.05 are bolded, and highlight when there is a positive association between the category of protein feature/functional site and a type of missense variants.
