## Supplementary Methods for "Mapping the Constrained Coding Regions in the human genome to their corresponding proteins"

*Aggregation of protein feature annotations and clinically interpreted variants*

*I. Protein features and functional annotations* (Figure 10, left blue-green panel) were mainly obtained from UniProtKB by querying the Proteins REST API (https://www.ebi.ac.uk/proteins/api/doc/index.html#/features). In particular, the annotations for residues involved in binding ligands: metals, small molecules, proteins, DNA/RNA and Cys forming disulphide bonds, were obtained and merged from both UniProtKB and VarSite (Laskowski et al. 2020), in March 2021. VarSite gathers its structural annotations via PDBsum (Laskowski et al. 2018) using structures from homologous proteins, and hence extending the information available for the amino acid positions. Additionally, we filtered out possible biologically non-relevant interactions by using BioLip (Yang, Roy, and Zhang 2013) artifact ligand list (https://zhanggroup.org/BioLiP/ligand_list). The feature “CATALYTIC” includes residues annotated as “ACT_SITE” in UniProtKB and/or as manually curated catalytic residues in M-CSA (Ribeiro et al. 2018) (https://www.ebi.ac.uk/thornton-srv/m-csa/api/residues/?format=json), accessed in June 2021. For ‘CDSjunctions’, we obtained the genomic coordinates of CDS (coding sequences) intron-exon boundaries from the Ensembl v101 GTF (Gene transfer format) file (http://ftp.ensembl.org/pub/release-101/gtf/homo_sapiens/Homo_sapiens.GRCh38.101.gtf.gz), and translated them to amino acid sequence following the methodology described in the “Mapping the CCRs to protein amino acids” section. “OTHER_REGION” and “OTHER_SITE” correspond to “Region” and “Site” of interest in the sequence annotations in UniProtKB, which cannot be described in other subsections of the sequence annotations. Amino acid conservation was obtained from VarSite, which uses Blastp to search for homologous sequences in UniProt, and then ScoreCons (Valdar 2002) for computing the conservation scores.

*II. Clinically interpreted missense variants* (Figure 10, middle yellow panel) were obtained from ClinVar (Landrum et al. 2018) (clinvar_20210814.vcf.gz, downloaded in August 2021 from https://ftp.ncbi.nlm.nih.gov/pub/clinvar/vcf_GRCh38/). We annotated the VCF file with Ensembl VEP v101 and restricted the dataset to those variants annotated as pathogenic/likely_pathogenic, benign/likely_benign, and combined the variants of uncertain significance (VUS) with those having conflicting interpretations of pathogenicity into a single subset named 'VUS_conflictive'.

*III. CCRs percentiles* (Figure 10, middle red panel) were obtained as described in the "*Mapping the CCRs to protein amino acids"* section.

*IV. Disorder and mobility related annotations* (Figure 10, right blue panel) were mainly obtained from the MobiDB database (Piovesan et al. 2021)(Mészáros et al. 2020) and DisProt (for manually curated intrinsically disordered regions) (Quaglia et al. 2022), assigning different levels of reliability. Here, we considered residue positions as disordered/mobile if they were true for any of the following annotations: "prediction_disorder_mobidb_lite", "curated_disorder_merge", "homology_disorder_merge", "prediction_disorder_th_50", "derived_missing_residues_context_dependent_th_90", "derived_missing_residues_th_90", "derived_mobile_context_dependent_th_90", "derived_mobile_th_90". Linear motifs were obtained from The Eukaryotic Linear Motif (ELM) resource (Kumar et al. 2020) (http://elm.eu.org/downloads.html#instances) in February 2021, requesting "true positives" and "Human" instances.

*Odds ratios tests for enrichment*

To calculate the OR, we compare the observed frequency with the enrichment of such CCRpct intervals with all other protein feature annotations. To do so, a 2x2 contingency table was constructed for each combination of feature annotation and CCRpct interval with the following cells: (a) the count of all residues at the CCRpct group that intersected the given feature annotation, (b) the count of all residues outside the CCRpct group that intersected the given feature annotation, (c) the count of all residues at the CCRpct group that intersect with other different feature annotations, and (d) the count of all residues outside the CCRpct group that intersect with other different feature annotations. The two-tailed Fisher's and one-tailed exact tests (Fisher 1970) were used to estimate the P-value and 95% confidence interval (CI 95%) for the OR of each contingency table, by testing the null hypothesis of a uniform distribution of CCRpct groups intersecting protein features and using *fisher.test* function from the *stats* library in R.

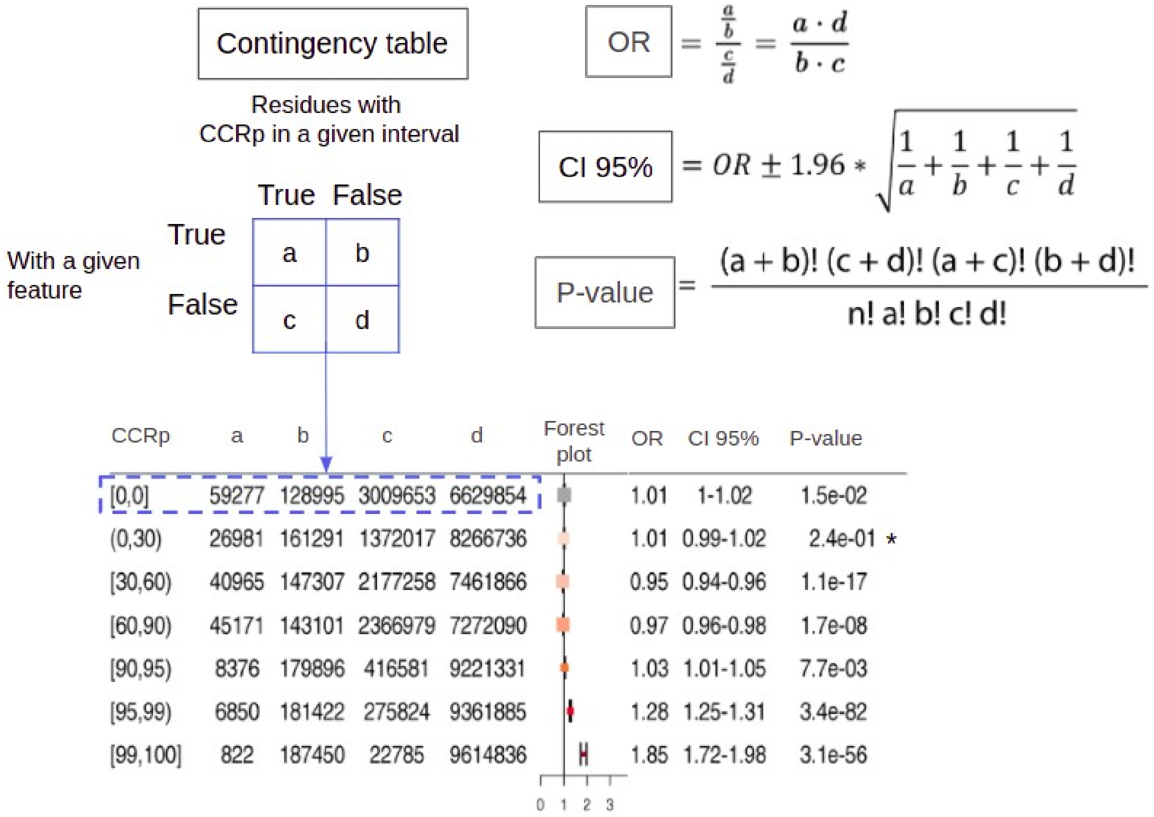


**Supplementary Methods Figure 1:** Example illustrating how 2x2 contingency tables were built for performing the odds ratios (OR) tests by counting residues presenting the co-ocurrence of the different CCRs percentiles groups with a protein feature, i.e. any of DOMAIN, CATALYTIC, DISORDER_MOBILE, etc or in NO_FEATURE for absence of any annotation or with the presence of PATHOGENIC, BENIGNG or VUS_conflict missense variants. From the OR test, three scenarios are possible: (1) if OR > 1.0 along with the CI 95% and P-value<=0.05: the residues with the given feature are OR times more likely associated to the given CCRp group, (2) OR < 1.0 along with the 95% CI and p-value<=0.05: the residues with the given feature are OR times less likely to have the given CCRp, and (3) the 95% confidence interval crosses over OR =1.0 and p-value>0.05, null hypothesis cannot be rejected hence no significant association is observed. A star (*) highlights this last situation.

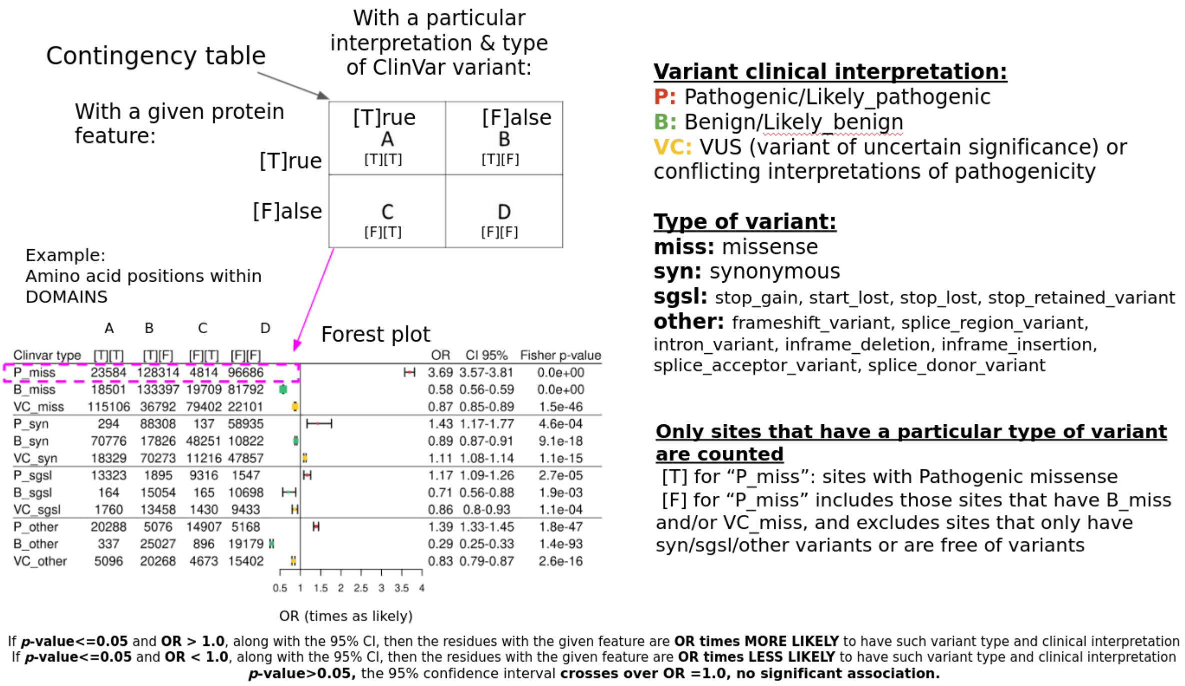


**Supplementary Methods Figure 2:** Example illustrating how contingency tables were built for performing the odds ratios (OR) tests, comparing each protein feature co-occurrence with different types of clinically interpreted variants from ClinVar.

### *References*


Fisher, Sir Ronald Aylmer. 1970. *Statistical Methods for Research Workers*. Oliver and Boyd.

Kumar, Manjeet, Marc Gouw, Sushama Michael, Hugo Sámano-Sánchez, Rita Pancsa, Juliana Glavina, Athina Diakogianni, et al. 2020. “ELM-the Eukaryotic Linear Motif Resource in 2020.” *Nucleic Acids Research* 48 (D1): D296–306.

Landrum, Melissa J., Jennifer M. Lee, Mark Benson, Garth R. Brown, Chen Chao, Shanmuga Chitipiralla, Baoshan Gu, et al. 2018. “ClinVar: Improving Access to Variant Interpretations and Supporting Evidence.” *Nucleic Acids Research* 46 (D1): D1062–67.

Laskowski, Roman A., Jagoda Jabłońska, Lukáš Pravda, Radka Svobodová Vařeková, and Janet M. Thornton. 2018. “PDBsum: Structural Summaries of PDB Entries.” *Protein Science: A Publication of the Protein Society* 27 (1): 129–34.

Laskowski, Roman A., James D. Stephenson, Ian Sillitoe, Christine A. Orengo, and Janet M. Thornton. 2020. “VarSite: Disease Variants and Protein Structure.” *Protein Science: A Publication of the Protein Society* 29 (1): 111–19.

Mészáros, Bálint, Gábor Erdős, Beáta Szabó, Éva Schád, Ágnes Tantos, Rawan Abukhairan, Tamás Horváth, et al. 2020. “PhaSePro: The Database of Proteins Driving Liquid-Liquid Phase Separation.” *Nucleic Acids Research* 48 (D1): D360–67.

Piovesan, Damiano, Marco Necci, Nahuel Escobedo, Alexander Miguel Monzon, András Hatos, Ivan Mičetić, Federica Quaglia, et al. 2021. “MobiDB: Intrinsically Disordered Proteins in 2021.” *Nucleic Acids Research* 49 (D1): D361–67.

Quaglia, Federica, Bálint Mészáros, Edoardo Salladini, András Hatos, Rita Pancsa, Lucía B. Chemes, Mátyás Pajkos, et al. 2022. “DisProt in 2022: Improved Quality and Accessibility of Protein Intrinsic Disorder Annotation.” *Nucleic Acids Research* 50 (D1): D480–87.

Ribeiro, António J. M., Gemma L. Holliday, Nicholas Furnham, Jonathan D. Tyzack, Katherine Ferris, and Janet M. Thornton. 2018. “Mechanism and Catalytic Site Atlas (M-CSA): A Database of Enzyme Reaction Mechanisms and Active Sites.” *Nucleic Acids Research* 46 (D1): D618–23.

Valdar, William S. J. 2002. “Scoring Residue Conservation.” *Proteins* 48 (2): 227–41.

Yang, Jianyi, Ambrish Roy, and Yang Zhang. 2013. “BioLiP: A Semi-Manually Curated Database for Biologically Relevant Ligand-Protein Interactions.” *Nucleic Acids Research* 41 (Database issue): D1096–1103.
